# Molecular analysis of 55 dopamine transporter variants from psychiatric patients reveals convergent mechanisms of dysfunction

**DOI:** 10.64898/2026.11.17.767405

**Authors:** Anna Campana, Leticia Alves da Silva, Jade Degrandmaison, Jonatan Fullerton Støier, Nina Kastner, Lucia Amalia Paggi, Fatemeh Kooti, Carmen Klein Herenbrink, Amy Hauck Newman, Harald H. Sitte, Thomas Werge, Freja Herborg, Thomas Stockner, Ulrik Gether

## Abstract

Rare genetic variants are increasingly recognized as contributors to neuropsychiatric disease and may reveal underlying pathobiological mechanisms. The dopamine transporter (DAT), encoded by SLC6A3, regulates dopaminergic neurotransmission through rapid dopamine (DA) reuptake. Although disease-associated DAT variants have been linked to disorders ranging from infantile parkinsonism to neuropsychiatric disease, the molecular basis of DAT dysfunction remains poorly understood. Here, we systematically investigated 55 DAT missense variants identified in the iPSYCH2012-nested case-control cohort. By assessing transport activity, inhibitor binding, cell-surface expression, subcellular localization, amphetamine responsiveness and constitutive DA efflux, we resolved the variants into seven phenotypic classes. These reflected distinct molecular defects, including impaired substrate and ion coupling, destabilized intramolecular interactions, and defective folding, trafficking or surface expression. Molecular dynamics simulations provided structural insight into representative variants. Collectively, our findings reveal mechanistic convergence of genetically diverse DAT variants and establish a framework for interpreting the pathogenicity of newly identified SLC6A3 variants.

## INTRODUCTION

Dopamine (DA) modulates multiple functions in the brain including reward-driven behaviors, motivation, learning and movement. Within dopaminergic circuits, the dopamine transporter (DAT) shapes extracellular DA tone by selective, sodium- and chloride-dependent reuptake of DA, thereby terminating DA signaling and replenishing presynaptic DA stores^1–3^. The important modulatory role of DA becomes apparent in pathological conditions involving dopaminergic dysregulation. Loss of dopaminergic neurons in the substantia nigra is a hallmark in Parkinson’s disease (PD), and extensive evidence implicates altered DA signaling in neuropsychiatric conditions, including – but not restricted to - attention deficit-hyperactivity disorder (ADHD), schizophrenia (SCZ), autism spectrum disorder (ASD), bipolar disorder (BD), depression and substance use disorder^2,4–7^.

A direct link between DAT and dopaminergic pathologies emerged by the discovery of inactivating genetic variants in the DAT gene (*DAT1 or SLC6A3*) causing the autosomal recessive “transportopathy”, DAT deficiency syndrome (DTDS), which manifests as infantile parkinsonism-dystonia with symptom onset within six months^8,9^. Subsequent studies, including studies from our lab, have shown that if residual uptake capacity is retained by the genetic variants, the disease may present as “atypical DTDS” characterized by childhood-onset neuropsychiatric disorder followed by adult-onset of parkinsonian symptoms^10–12^. In parallel, an increasing number of rare or ultra-rare DAT missense mutants have been identified in individuals diagnosed with neuropsychiatric conditions such as ASD, BD and ADHD^12–18^. The rather few variants characterized so far display different functional phenotypes, including variable degrees of hypofunction and distinct gain-of-function effects^12,14–16,19^. One interesting gain of function phenotype is constitutive efflux of DA via DAT, which was identified in variants such as T356M, which has been linked to ADHD and autism^14–16,19^; however, the prevalence, structural basis and pathophysiological significance of this phenotype remain unclear. Together, the available data suggest a possible gradient in disorder onset and severity linked to the significance of the molecular phenotype associated with genetic DAT variants^13,14,20^. Nevertheless, only a limited number of variants have been functionally investigated, leaving a major knowledge gap on how molecular alterations in DAT can have disease-related effects.

Although rare and ultra-rare coding variants account for only a fraction of genetic risk, individual variants can exert large biological effects and provide uniquely informative entry points into disease mechanisms. Indeed, growing evidence supports an important contribution of rare variation to neuropsychiatric disorders^21^. Therefore, to improve our understanding of how specific alterations in DAT function translate into clinical phenotypes, we conducted a systematic analysis of rare, patient-derived missense genetic variants in the *SLC6A3* gene. This effort leveraged whole exome sequencing (WES) data obtained from the iPSYCH2012 case-cohort study, which included individuals with confirmed diagnosis of at least one of these five disorders: ADHD, ASD, BD, single and recurrent depression (SaRD), and SCZ^22^. We identified 83 SLC6A3 missense variants, of which 55 were prioritized for detailed study based on their enrichment within disease groups (>50%). By integrating analyses of substrate uptake, inhibitor binding, expression, constitutive and amphetamine-induced DA efflux, and molecular dynamics simulations, we resolved these variants into seven phenotypic categories representing distinct molecular modes of DAT dysfunction. Notably, the variants with complete or almost complete loss of function were selectively identified among cases, supporting association between severe functional impairment and disease phenotype. In summary, our results uncover a broad yet convergent landscape of DAT dysfunction arising from disease linked DAT variants and establish a foundation for interpreting the pathogenicity of *SLC6A3* variation in neuropsychiatric disorders.

## RESULTS

### Selection of DAT missense variants identified in neuropsychiatric individuals

To delineate the contribution of DAT dysfunction to psychiatric manifestations, we employed large-scale WES data from the iPSYCH2012-nested case-control study^22^. The study encompassed 19,851 individuals and identified 83 missense variants in *SLC6A3*, of which 55 were predominantly represented in the disease group and selected for detailed analysis (Fig. 1a, b and Table 1).

**Figure 1.**
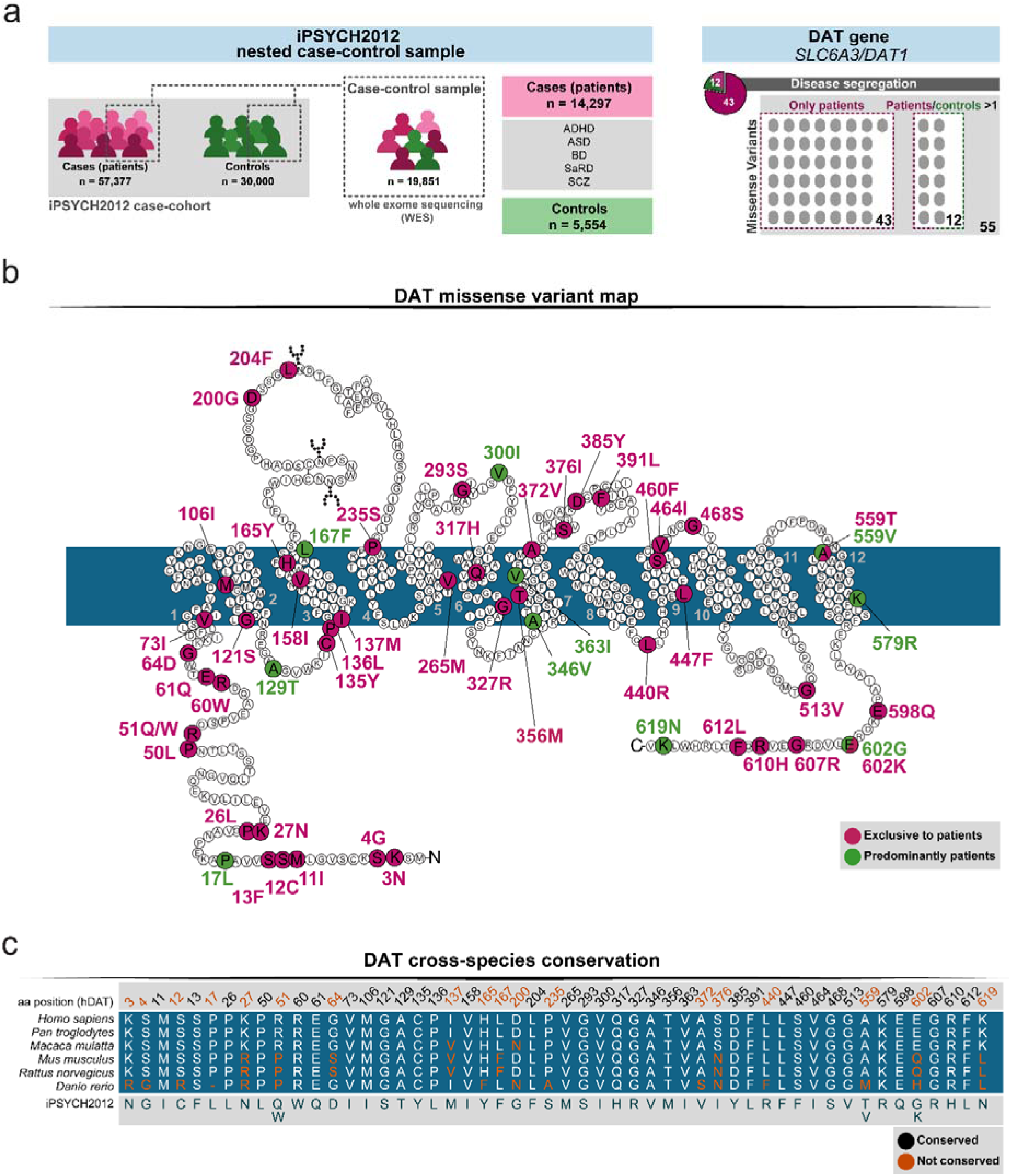
Schematic overview of the study, snake diagram of human DAT mapping missense mutants selected for functional and structural characterization, and sequence alignment of DAT orthologues. **a** Workflow of the study including source of genetic information, subsequent selection, and disease segregation of the DAT variants selected for this study. **b** Illustration of the DAT amino acid sequence and membrane topology. Missense mutants identified in the iPSYCH2012 cohort are highlighted by enlarged circles; variants exclusively detected in the case group are shown in magenta, whereas variants identified in both patients and controls are shown in green. Residues A559 and E602 each harbor two distinct mutations in the iPSYCH2012 cohort, and are thus depicted in both magenta and green, according to their complete or partial segregation to the disease group. **c** Sequence alignment of human DAT (*Homo sapiens*) residues mutated in this study with orthologue DAT sequences from *Pan troglodytes*, *Macaca mulatta*, *Mus musculus*, *Rattus norvegicus* and *Danio rerio*.

**Table 1.** iPSYCH2012 diagnostic information, ClinVar annotations and predicted impairment scores of DAT missense variants.

|  | iPSYCH2012 |  |  | ClinVar |  | gnomAD |  | PolyPhen-2 |
|---|---|---|---|---|---|---|---|---|
| Mutation | P | C | Phenotype | Phenotype | n | CADD (PHRED) score | AF (‰) | Score |
| K3N | yes | no | ASD | - | - | 9.75 | 0.007434 | 0.047 (B) |
| S4G | yes | no | SCZ/ SCZ, ADHD, ASD | - | - | - | - | 0.00 (B) |
| M11I | yes | no | SCZ | IGD (1) / IPD (1) / np (1) | 3 | 20.8 | 0.003717 (c.33G>A) | 0.031 (B) |
| S12C | yes | no | SCZ | - | - | 24.9 | 0.003098 | 0.984 (D) |
| S13F | yes | no | BD | - | - | - | - | 0.993 (D) |
| P17L | yes | yes | ASD / BD | - | - | 21.4 | 0.01487 | 0.014 (B) |
| P26L | yes | no | ASD | IGD (1) / IPD (1) / np (1) | 3 | 20.6 | 0.04956 | 0.900 (P) |
| K27N | yes | no | BD | IGD (1) / IPD (1) | 2 | 23.1 | 0.006814 | 0.783 (P) |
| P50L | yes | yes | ASD / SCZ | DTDS, susceptibility to tobacco addiction (1) / IPD (1) | 2 | 18.2 | 0.05080 | 0.8 (P) |
| R51Q | yes | no | ASD / ADHD | IGD (1) / IPD (1) | 2 | 9.81 | 0.03717 | 0.00 (B) |
| R51W | yes | no | ADHD, SCZ | IPD | 1 | 15.6 | 0.03097 | 0.536 (P) |
| R60W | yes | no | ADHD, BD, SaRD | DTDS (1) / IPD (1) | 2 | 28.2 | 0.001239 | 1.00 (D) |
| E61Q | yes | no | ADHD | - | - | - | - | 0.999 (D) |
| G64D | yes | no | ADHD / SCZ | - | - | - | - | 0.007 (B) |
| V73I | yes | no | ADHD / ASD | DTDS | 1 | 23.5 | 0.03346 | 0.774 (P) |
| M106I | yes | no | SCZ | - | - | - | - | 0.152 (B) |
| G121S | yes | no | ADHD / BD | - | - | 25.7 | 0.003717 | 0.998 (D) |
| A129T | yes | yes | ADHD / ADHD, SCZ, SaRD | IPD | 1 | 23.8 | 0.1140 | 0.993 (D) |
| C135Y | yes | no | ASD | - | - | - | - | 1.00 (D) |
| P136L | yes | no | SCZ | - | - | - | - | 0.998 (D) |
| I137M | yes | no | ADHD | DTDS | 1 | 18.6 | 0.008054 | 0.248 (B) |
| V158I | yes | no | BD | IGD (1) / IPD (1) | 2 | 24.2 | 0.005576 | 0.995 (D) |
| H165Y | yes | no | SCZ | IPD | 1 | 18.9 | 0.001859 | 0.00 (B) |
| L167F | 63 | yes | ADHD / ADHD, ASD / ADHD, ASD, BD / ADHD, BD / ADHD, BD, SaRD / ADHD, SaRD / ADHD, SCZ / ADHD, SCZ, SaRD / ASD / ASD, SaRD / ASD, SCZ / ASD, SCZ, SaRD/ BD / BD, SaRD / BD, ASD / BD, ASD, SaRD / BD, SCZ, SaRD / SCZ / SCZ, SaRD | DTDS (2), of which (1) susceptibility to tobacco addiction / IPD (1) / np (5) | 8 | 13.8 | 1.035 | 0.036 (B) |
| D200G | yes | no | BD / SCZ | - | - | 4.63 | 0.001239 | 0.00 (B) |
| L204F | yes | no | SCZ | - | - | 4.33 | 0.001239 | 0.001 (B) |
| P235S | yes | no | ADHD | - | - | 13.4 | 0.001240 | 0.003 (B) |
| V265M | yes | no | ADHD | - | - | - | - | 1.00 (D) |
| G293S | yes | no | ADHD / ADHD, BD | - | - | 24.3 | 0.0006197 | 0.900 (D) |
| V300I | 12 | yes | ADHD / ADHD, ASD / ASD / ASD, SCZ / BD / BD, SaRD / SCZ, SaRD | DTDS, susceptibility to tobacco addiction (1) / IPD (1) / np (1) | 3 | 11.8 | 0.1723 | 0.027 (B) |
| Q317H | yes | no | ASD | - | - | - | - | 1.00 (D) |
| G327R | yes | no | ADHD | - | - | 26.9 | 0.002479 | 1.00 (D) |
| A346V | yes | yes | ADHD / ASD | IPD | 1 | 25.1 | 0.07564 | 1.00 (D) |
| T356M | yes | no | ADHD, SaRD / ASD | DTDS (1) / IPD (1) | 2 | 28.8 | 0.02542 | 1.00 (D) |
| V363I | yes | yes | ASD | IPD | 1 | 25.4 | 0.01426 | 0.941 (D) |
| A372V | yes | no | ADHD | - | - | - | - | 0.971 (D) |
| S376I | yes | no | ADHD | - | - | - | - | 0.248 (B) |
| D385Y | yes | no | ADHD / ADHD, ASD | - | - | 26.5 | 0.0006202 | 0.999 (D) |
| F391L | yes | no | ADHD / ASD | - | - | - | - | 0.441 (B) |
| L440R | yes | no | ADHD / SCZ | - | - | 22.8 | 0.009917 | 0.006 (B) |
| L447F | yes | no | ADHD, SCZ | - | - | 24.9 | 0.006199 | 0.696 (P) |
| S460F | yes | no | ASD | - | - | - | - | 0.980 (D) |
| V464I | yes | no | ASD | IPD (1) / np (2) | 3 | 20.8 | 0.2275 | 0.836 (P) |
| G468S | yes | no | ADHD / ASD | - | - | 29.1 | 0.0006220 | 1.00 (D) |
| G513V | yes | no | BD | - | - | - | - | 1.00 (D) |
| A559T | yes | no | ADHD, ASD | IPD | 1 | 16.3 | 0.02045 | 0.003 (B) |
| A559V | 35 | yes | ADHD / ADHD, ASD / ADHD, BD, SCZ, SaRD / ADHD, SaRD / ADHD, SCZ / ASD / ASD, SaRD / ASD, SCZ, SaRD / BD / SCZ / SCZ, SaRD | DTDS (1) / IGD (1) / IPD (1) / np (3) / other (1) | 7 | 7.97 | 1.227 | 0.00 (B) |
| K579R | yes | yes | ADHD / ADHD, ASD / ASD / SCZ | IPD (1) / np (1) | 2 | 24.8 | 0.09543 | 0.324 (B) |
| E598Q | yes | no | ADHD | - | - | 22.7 | 0.001252 | 0.889 (D) |
| E602G | 8 | yes | ADHD / ASD / BD / SCZ | IGD (1) / IPD (1) / np (3) | 5 | 14.1 | 0.4823 | 0.00 (B) |
| E602K | yes | no | BD, SaRD | - | - | - | - | 0.021 (B) |
| G607R | yes | no | SCZ | - | - | - | - | 0.136 (B) |
| R610H | yes | yes | ADHD / ASD, SaRD | IPD (1) / np (1) | 2 | 26.5 | 0.03466 | 0.993 (D) |
| F612L | yes | no | ASD | - | - | 24.3 | 0.0006313 | 0.721 (P) |
| K619N | 21 | yes | ADHD / ADHD, ASD / ADHD, BD / ASD / BD / BD, SCZ / SCZ | IGD (1) / IPD (1) / np (2) | 4 | 16.1 | 0.3129 | 0.003 (B) |

For each missense variant, the table reports the exact number of patients (P, second column), when more than five patients were identified, or "yes" when fewer than five patients were identified as carrying the respective mutation in the iPSYCH2012 dataset. Along with correspondent disease diagnosis. For each mutation, the absence or presence of control individuals (C, third column) is reported as "no" or "yes", respectively. iPSYCH2012 mutants registered on the ClinVar database are annotated together with their associated disease conditions. For iPSYCH2012 variants annotated on gnomAD, CADD-PHRED scores indicate predicted deleteriousness relative to all possible variants in the human genome. The population allele frequency (AF, ‰) is also reported as provided by gnomAD. The predicted damage severity imposed by each mutation was assessed using the PolyPhen-2 tool and assigned with a score ranging from 0 to 1. ADHD, attention-deficit/hyperactivity disorder; ASD, autism spectrum disorder; BD, bipolar disorder; DTDS, dopamine transporter deficiency syndrome; IGD, inborn genetic diseases; IPD, infantile parkinsonism-dystonia; np, not provided; SCZ, schizophrenia; SaRD, single and recurrent depression. B = "benign"; P (last column) = "possibly damaging"; D = "probably damaging" or "damaging". P (second column), patients; C, controls; n, number of entries; CADD, Combined Annotation Dependent Depletion; gnomAD, Genome Aggregation Database.

We first interrogated the ClinVar database and found that 24 out of 55 variants (43.6%) had been previously reported, whereas 31 variants (56.4%) were novel to our study (Table 1). When assessing the potential impact of each amino acid substitution, we found 31 variants scoring as damaging or possibly damaging according to the Polymorphism Phenotyping v2 (PolyPhen-2) software tool (Table 1). We additionally report allele frequency and predicted deleteriousness according to the combined annotation dependent depletion (CADD) tool, as annotated in the Genome Aggregation Database (gnomAD) (Table 1).

### Distinct perturbations cause kinetic impairment in DAT disease mutants

To dissect molecular phenotypes of the selected DAT coding variants, we first tested DA transport activity using a cell-based radioligand uptake assay (Fig. 2, p*K*_m_ (−log*K*_m_ [M]) and *V*_max_ values reported in Table 2). Second, we performed a binding assay using a fluorescently tagged, membrane-impermeable cocaine analogue (DG3-80) (Fig. 3, p*K*_d_ (−log*K*_d_ [M]) and *B*_max_ values reported in Table 2, Scatchard plots reported in Supplementary Fig. 1)^23^.

**Figure 2.**
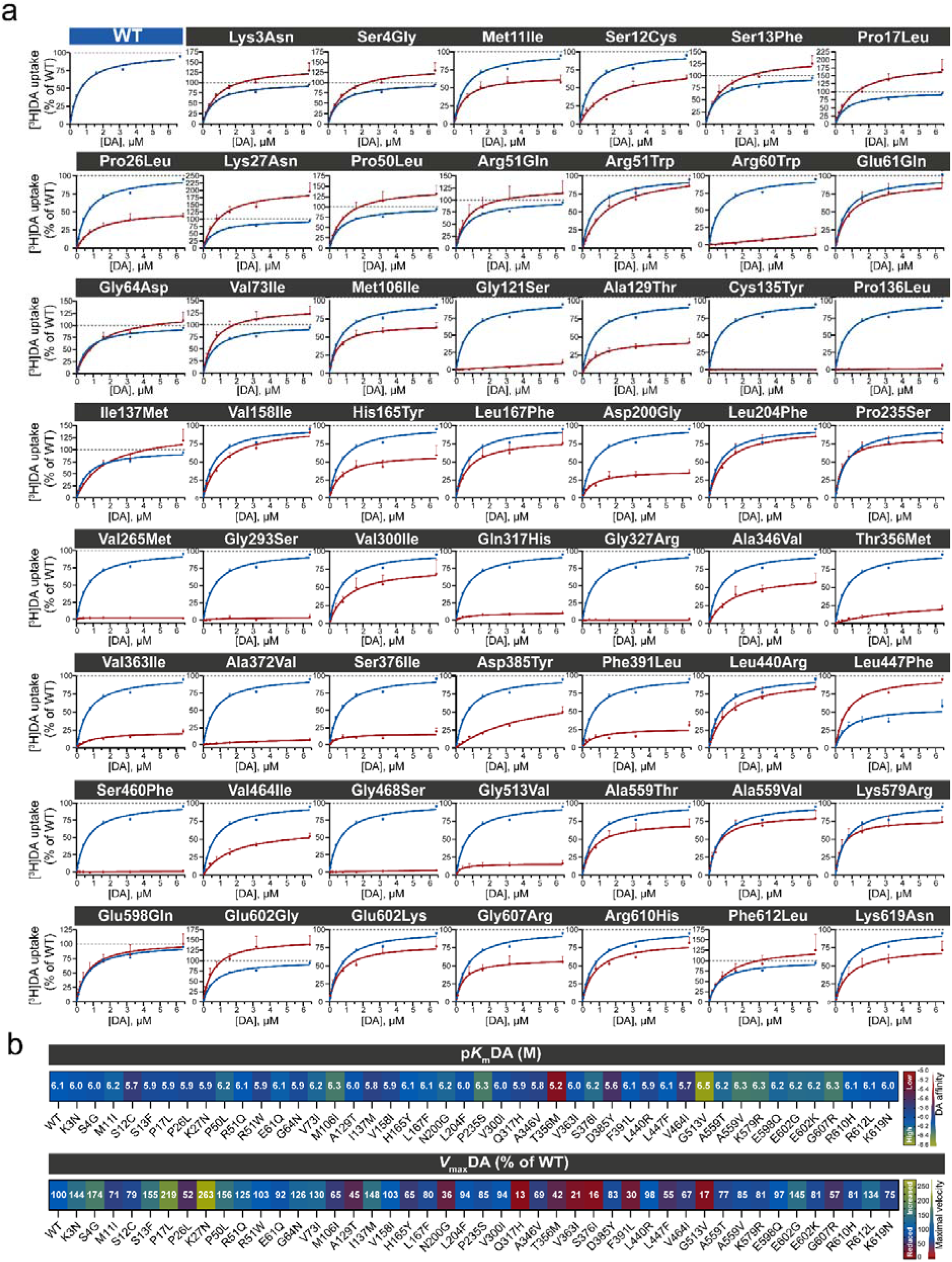
[^3^H]DA saturation uptake reveals different degrees of kinetic impairment in DAT disease variants. **a** Systematic assessment of DA uptake capacity of each DAT mutant through [^3^H]DA saturation uptake in transiently transfected HEK293 cells. All curves represent the average ± S.E.M. of 3 to 8 independent experiments, each conducted in duplicates, and normalized to the maximal velocity of the WT. Mutants (red) are displayed in direct comparison to the WT (blue). **b** Heat maps of apparent DA affinity (DA K_m_, upper row) and maximal transport velocity (DA V_max_, lower row) calculated by fitting the values in panel **a** using the Michaelis-Menten equation. DAT variants for which DA uptake was not detected, and kinetic values could not be fitted, are excluded from panel b.

**Figure 3.**
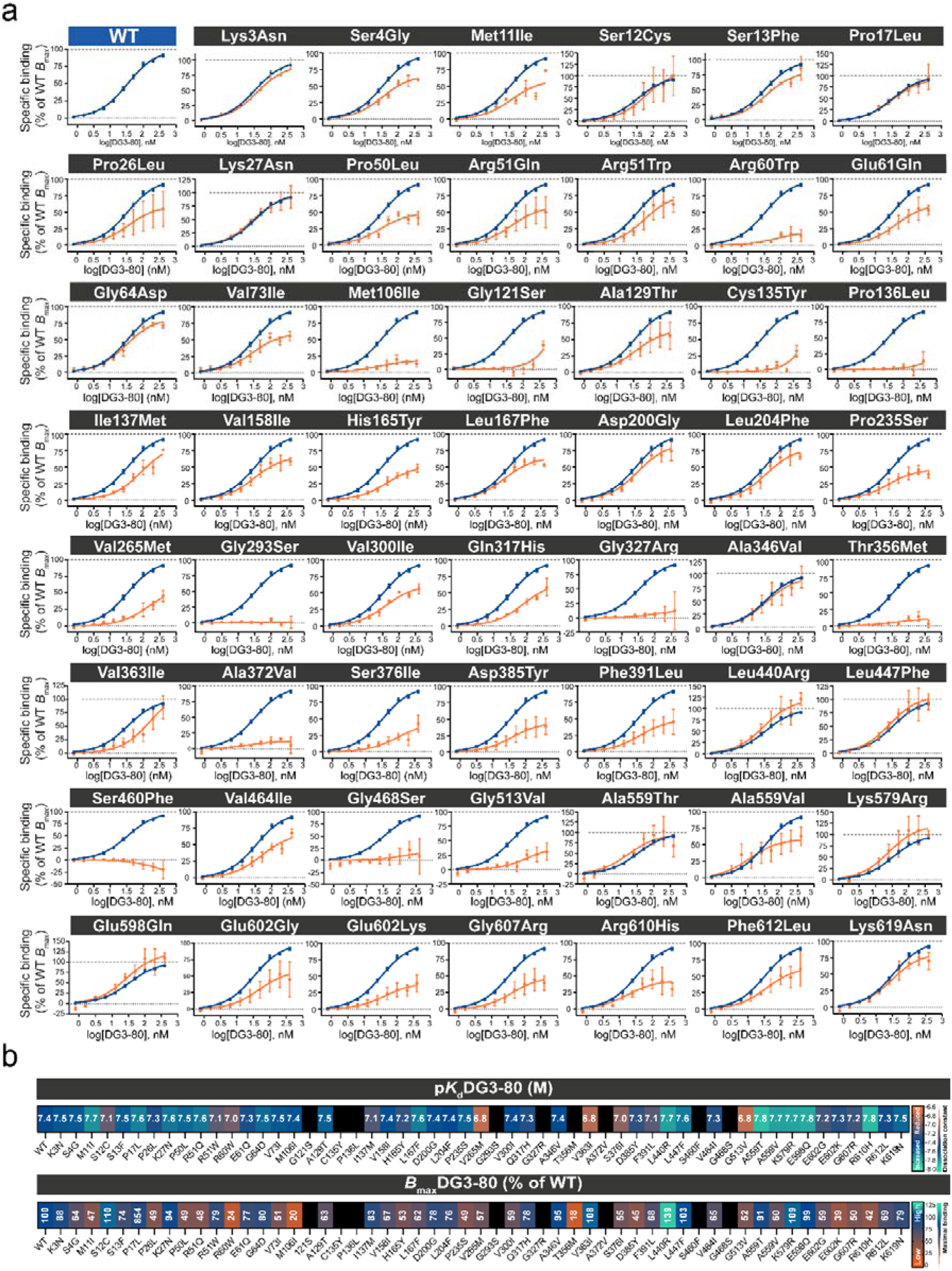
Neuropsychiatric DAT variants display altered binding properties. **a** Saturation binding of fluorescently conjugated cocaine analogue DG3-80, performed on transiently transfected HEK293 cells expressing the WT (blue) or single DAT coding variants (orange). Binding curves were fitted using a one-site specific binding nonlinear regression analysis on baseline-subtracted data. Single data points represent mean ± S.E.M. of at least three independent experiments, each performed in duplicates and normalized to apparent maximal binding of the WT. **b** Heat maps of DG3-80 dissociation constant (DG3-80 *K*_d_, upper row) and maximal binding (DG3-80 *B*_max_, lower row) obtained from fitted curves shown in panel **a**. DAT mutants with undetectable DG3-80 binding and therefore no fitted dissociation values, are displayed as black squares.

**Table 2.** Kinetic values calculated from saturation [^3^H]DA uptake and saturation DG3-80 binding.

|  | p*K*<sub>m</sub>DA, mean ± S.E.M. (M) | *V*<sub>max</sub>DA, mean ± S.E.M. (% of WT) | n (DA uptake) | p*K*<sub>d</sub>DG3-80, mean ± S.E.M. (M) | *B*<sub>max</sub> DG3-80, mean ± S.E.M. (% of WT) | n (DG3-80 binding) |
|---|---|---|---|---|---|---|
| WT | 6.10 ± 0.0 | 100.0 | 41 | 7.39 ± 0.1 | 100.0 | 23 |
| Lys3Asn | 6.05 ± 0.2 | 144 ± 31 | 5 | 7.46 ± 0.1 | 88 ± 14 | 3 |
| Ser4Gly | 6.01 ± 0.1 | 174 ± 48<sup>*</sup> | 5 | 7.47 ± 0.0 | 64 ± 4 | 3 |
| Met11Ile | 6.15 ± 0.1 | 71 ± 12 | 5 | 7.67 ± 0.2 | 47 ± 16 | 3 |
| Ser12Cys | 5.74 ± 0.1 | 79 ± 7 | 4 | 7.08 ± 0.2 | 110 ± 42 | 3 |
| Ser13Phe | 5.92 ± 0.2 | 155 ± 25 | 5 | 7.48 ± 0.2 | 74 ± 17 | 3 |
| Pro17Leu | 5.95 ± 0.2 | 219± 78<sup>****</sup> | 4 | 7.61 ± 0.1 | 85 ± 23 | 4 |
| Pro26Leu | 5.90 ± 0.0 | 52 ± 7 | 4 | 7.31 ± 0.2 | 49 ± 23<sup>*</sup> | 4 |
| Lys27Asn | 5.90 ± 0.2 | 263 ± 96<sup>****</sup> | 4 | 7.58 ± 0.1 | 94 ± 20 | 3 |
| Pro50Leu | 6.21 ± 0.1 | 156 ± 30 | 5 | 7.52 ± 0.0 | 49 ± 8 | 3 |
| Arg51Gln | 6.07 ± 0.1 | 125 ± 26 | 6 | 7.58 ± 0.0 | 48 ± 20 | 3 |
| Arg51Trp | 5.85 ± 0.1 | 103 ± 18 | 4 | 7.11 ± 0.2 | 79 ± 15 | 3 |
| Arg60Trp | ND | Not fitted | 6 | 7.03 ± 0.2 | 24 ± 12<sup>****</sup> | 4 |
| Glu61Gln | 6.13 ± 0.1 | 92 ± 19 | 5 | 7.32 ± 0.1 | 77 ± 19 | 4 |
| Gly64Asp | 5.92 ± 0.1 | 126 ± 20 | 5 | 7.53 ± 0.2 | 80 ± 5 | 3 |
| Val73Ile | 6.19 ± 0.1 | 130 ± 18 | 5 | 7.46 ± 0.2 | 51 ± 5 | 3 |
| Met106Ile | 6.31 ± 0.1 | 65 ± 7 | 5 | 7.44 ± 0.1 | 18 ± 5<sup>****</sup> | 4 |
| Gly121Ser | ND | Not fitted | 4 | ND | Not fitted | 4 |
| Ala129Thr | 6.03 ± 0.0 | 45 ± 6 | 4 | 7.48 ± 0.2 | 63 ± 23 | 3 |
| Cys135Tyr | ND | Not fitted | 4 | ND | Not fitted | 4 |
| Pro136Leu | ND | Not fitted | 3 | ND | Not fitted | 4 |
| Ile137Met | 5.81 ± 0.1 | 148 ± 42 | 5 | 7.14 ± 0.1 | 72 ± 11 | 4 |
| Val158Ile | 5.88 ± 0.1 | 103 ± 10 | 5 | 7.43 ± 0.2 | 67 ± 3 | 3 |
| His165Tyr | 6.11 ± 0.2 | 65 ± 15 | 6 | 7.23 ± 0.1 | 53 ± 3 | 3 |
| Leu167Phe | 6.13 ± 0.1 | 80 ± 19 | 6 | 7.60 ± 0.0 | 62 ± 4 | 3 |
| Asp200Gly | 6.19 ± 0.1 | 36 ± 6 | 5 | 7.43 ± 0.1 | 73 ± 13 | 3 |
| Leu204Phe | 6.01 ± 0.1 | 94 ± 11 | 5 | 7.40 ± 0.1 | 76 ± 9 | 3 |
| Pro235Ser | 6.31 ± 0.1 | 85 ± 10 | 5 | 7.51 ± 0.2 | 49 ± 6 | 3 |
| Val265Met | ND | Not fitted | 3 | 6.80 ± 0.1 | 57 ± 7 | 3 |
| Gly293Ser | ND | Not fitted | 3 | ND | Not fitted | 3 |
| Val300Ile | 5.96 ± 0.2 | 94 ± 31 | 6 | 7.39 ± 0.0 | 59 ± 5 | 3 |
| Gln317His | 5.87 ± 0.2 | 13 ± 4<sup>***</sup> | 6 | 7.30 ± 0.1 | 54 ± 6 | 4 |
| Gly327Arg | ND | Not fitted | 5 | ND | Not fitted | 3 |
| Ala346Val | 5.82 ± 0.1 | 69 ± 14 | 8 | 7.42 ± 0.2 | 95 ± 20 | 3 |
| Thr356Met | 5.20 ± 0.2<sup>****</sup> | 42 ± 7 | 3 | ND | Not fitted | 4 |
| Val363Ile | 6.01 ± 0.2 | 21 ± 3<sup>*</sup> | 6 | 6.84 ± 0.2 | 108 ± 18 | 3 |
| Ala372Val | ND | Not fitted | 5 | ND | Not fitted | 3 |
| Ser376Ile | 6.21 ± 0.3 | 16 ± 6<sup>**</sup> | 4 | 6.96 ± 0.4 | 55 ± 21 | 3 |
| Asp385Tyr | 5.62 ± 0.3 | 83 ± 21 | 4 | 7.25 ± 0.2 | 45 ± 13 | 3 |
| Phe391Leu | 6.06 ± 0.3 | 30 ± 5 | 4 | 7.12 ± 0.3 | 81 ± 27 | 3 |
| Leu440Arg | 5.88 ± 0.1 | 98 ± 12 | 5 | 7.73 ± 0.2 | 106 ± 28 | 4 |
| Leu447Phe | 6.11 ± 0.1 | 55 ± 9 | 4 | 7.59 ± 0.1 | 103 ± 23 | 3 |
| Ser460Phe | ND | Not fitted | 3 | ND | Not fitted | 3 |
| Val464Ile | 5.72 ± 0.2 | 67 ± 7 | 3 | 7.29 ± 0.2 | 65 ± 2 | 3 |
| Gly468Ser | ND | Not fitted | 3 | ND | Not fitted | 4 |
| Gly513Val | 6.54 ± 0.1 | 17 ± 5<sup>*</sup> | 4 | 6.79 ± 0.4 | 101 ± 66 | 3 |
| Ala559Thr | 6.20 ± 0.1 | 77 ± 11 | 6 | 7.83 ± 0.1 | 88 ± 29 | 3 |
| Ala559Val | 6.31 ± 0.1 | 85 ± 9 | 5 | 7.72 ± 0.4 | 60 ± 9 | 3 |
| Lys579Arg | 6.30 ± 0.1 | 81 ± 13 | 6 | 7.74 ± 0.2 | 109 ± 27 | 3 |
| Glu598Gln | 6.18 ± 0.1 | 97 ± 15 | 5 | 7.76 ± 0.1 | 99 ± 26 | 3 |
| Glu602Gly | 6.22 ± 0.1 | 145 ± 21 | 4 | 7.19 ± 0.2 | 65 ± 14 | 3 |
| Glu602Lys | 6.21 ± 0.2 | 81 ± 12 | 5 | 7.27 ± 0.2 | 42 ± 7<sup>*</sup> | 3 |
| Gly607Arg | 6.31 ± 0.1 | 57 ± 9 | 3 | 7.20 ± 0.3 | 55 ± 9 | 3 |
| Arg610His | 6.11 ± 0.1 | 81 ± 5 | 3 | 7.82 ± 0.1 | 42 ± 15<sup>*</sup> | 3 |
| Phe612Leu | 6.05 ± 0.2 | 134 ± 49 | 3 | 7.34 ± 0.3 | 69 ± 19 | 3 |
| Lys619Asn | 6.03 ± 0.2 | 75 ± 11 | 4 | 7.52 ± 0.1 | 79 ± 11 | 3 |

Data are p*K*<sub>m</sub> (–log*K*<sub>m</sub> [M]) and *V*<sub>max</sub> values for DA calculated from DA uptake experiments using the Michaelis-Menten equation. p*K*<sub>d</sub> (–log*K*<sub>d</sub> [M]) and *B*<sub>max</sub> values for DG3-80 were obtained using a one-site specific binding nonlinear regression analysis of DG3-80 binding curves. *V*<sub>max</sub> and *B*<sub>max</sub> values were normalized to the maximal velocity and maximum binding capacity of the WT, respectively. Data are mean ± S.E.M.; the number of biological replicates is reported in the table (n). For p*K*<sub>m</sub>DA, *V*<sub>max</sub>DA, p*K*<sub>d</sub>DG3-80 and *B*<sub>max</sub>DG3-80, statistical significance was evaluated using one-way ANOVA with Holm-Šídák′ multiple comparisons test (<sup>\*</sup>, *p* < 0.05; <sup>\*\*</sup>, *p* < 0.01 <sup>\*\*\*</sup>, *p* < 0.001; <sup>\*\*\*\*</sup>, *p* < 0.0001).

Based on the data from these two assays, we classified the mutations into seven groups:

Group 1: G121S, C135Y, P136L, G293S, G327R, A372V, S460F, G468S. For C135Y, P136L, G293S, G327R, S460F and G468S, the uptake assay yielded no detectable signal. For G121S and A372V, the maximal velocity was too low to reliably calculate p*K*_m_ values (Fig. 2 and Table 2). The marked loss of functional activity in these 8 variants might not be surprising given their location in the transporter (Fig.1b). G121S is situated on the intracellular edge of transmembrane segment 2 (TM2) close to TM6b, which is directly involved in substrate binding and translocation^24^. C135Y and P136L are located between intracellular loop 1 (ICL1) and TM3; TM3 participates in substrate binding while ICL1 establishes hydrogen bonds and electrostatic interactions with the C-terminus^24,25^. G293S is located on ECL3, proximal to TM7 and to the proposed DAT allosteric site, where ECL3 has been implicated in DAT biosynthesis and trafficking, and in substrate recognition in the homologous norepinephrine transporter (NET)^26–28^. G327R maps to TM6, which directly participates in substrate and ion binding^24,29^. A372V is located at the upper edge of TM7, which is part of the bundle domain^24,30^. Finally, S460F and G468S are found in TM 9 and 10, respectively, which are part of the scaffold domain of the transporter, with TM9 participating in DAT dimerization and oligomerization^30,31^.

In the DG3-80 binding assay, we observed no detectable binding for all the variants (G121S, C135Y, P136L, G293S, G327R, A372V, S460F, G468S) (Fig. 3 and Table 2), suggesting that the loss of DA uptake could be the result of either lack of surface expression of the mutated transporter or direct perturbation of substrate binding/transport and binding of cocaine-like inhibitors^27,28^. Of note, all 8 variants in this group, except A372, show high cross-species conservation (Fig. 1c) and are highly damaging according to PolyPhen-2 (Table 1). Remarkably, none of the mutants has previous ClinVar entry; however, all of them are solely found in the disease group pertaining to the iPSYCH2012 sample (Table 1).

Group 2: R60W and V265M. These mutants display loss of measurable DA uptake (Fig. 2 and Table 2) but partially preserved DG3-80 binding with significant reduction in maximal DG3-80 binding for R60W (Figs 2-3 and Table 2). This suggests that mainly catalytic impairment underlies the loss of uptake. For R60W, the phenotype is consistent with the well-documented role of R60 in stabilizing the intracellular gate of the transporter via formation of a salt bridge with D436 that is stabilized by a cation-π interaction between R60 and Y335^32,33^. Similarly, V265 locates to the inner face of TM5 and thus to a region proposed to regulate Na2 release and to influence the transition to the inward-open state^24,28,34^.

Group 3: A129T, D200G, Q317H, V363I, S376I, F391L and G513V. For Q317H, V363I, S376I, and G513V, we observed significantly lowered *V*_max_ for uptake (∼13%, ∼20%, ∼16% and ∼17% of WT, respectively) alongside unchanged p*K*_m_, For A129T, D200G and F391L, we also observed reduced *V*_max_ (∼30-45% of WT) but the values did not show up as significant (Fig. 2 and Table 2). For all mutants in the group, there were no significant changes in DG3-80 binding, but trends towards lower *B*_max_ were seen for Q317H and S376I. (Figs 2-3 and Table 2). The residues occupy various locations in the transporter: A129T, D200G, F391L and G513V are situated far away from the substrate binding site, on either extracellular- or intracellular-facing regions of DAT (Fig. 1b, c), suggesting an indirect effect on DA uptake capacity or surface targeting. Q317, however, resides on TM6 and coordinates chloride ion binding^24,35^. V363I is in a conserved and critical position located at the top of TM7, facing TM5 and indirectly affecting Na2 coordination (Fig. 1b, c)^24^. S376I is located to EL4, a region implicated in extracellular vestibule closure and control of substrate access to the S1 site (Fig. 1b, c)^28^. ^24^

Group 4: T356M was the only mutant that displayed significantly reduced DA affinity with no significant alteration of *V*_max_ combined with impaired DG3-80 binding (T356M, not fitted) (Figs. 2-3 and Table 2). T356M is found in TM7 facing TM5 and TM1, with likely implications for ion binding and substrate binding/transport^24,34^. Notably, T356M, which is strictly conserved (Fig. 1c), has previously been described and associated with ASD^14,16^.

Group 5: P26L, M106I, E602K and R610H showed significantly reduced *B*_max_ for DG3-80, with no significant changes in DA uptake profiles (Figs 2-3 and Table 2). However, there was a strong trend towards lower *V*_max_ for P26L (∼52% of WT), a conserved residue found in the N-terminus (Fig. 1b, c). M106 locates to TM2, which is part of the bundle domain participating in the outward-to-inward transitions during the transport cycle (Fig.1b)^28,30^. E602K and R610H, both C-terminal variants, result in the loss of a negatively and positively charged side chain, respectively (Fig.1b).

Group 6: S4G, P17L and K27N. These three N-terminal variants showed a significant increase in *V*_max_, while retaining WT-like p*K*_m_ and DG3-80 binding profiles (Figs 2-3 and Table 2). Interestingly, S4 is one out of five serines (S2, S4, S7, S12 and S13) that are clustered in the DAT N-terminus and constitute phosphorylation sites for protein kinase C - and Ca^2+^/calmodulin-dependent protein kinase II. Also, K27 is one of the three lysines in DAT subjected to ubiquitination^36–39^.

Group 7: K3N, M11I, S12C, S13F, P50L, R51Q, R51W, E61Q, G64D, V73I, I137M, V158I, H165Y, L167F, L204F, P235S, V300I, A346V, D385Y, L440R, L447F, V464I, A559T, A559V, K579R, E598Q, E602G, G607R, F612L and K619N. These variants display no significant alterations either in DA uptake profiles or DG3-80 binding (Figs 2-3 and Table 2). Within this sub-set of variants, L167F, V300I, A559V, K579R and K619N showed the lowest patients/(patients+controls) ratio (Table 1).

Although the selection of variants for functional analysis was inherently biased toward those preferentially observed in cases, we nevertheless examined the distribution of patients and controls across the functional groups. Because the protocol does not permit reporting the number of carriers when the total count is <5, cases could not be stratified by diagnosis and we had to split the seven groups into three broader categories: Groups 1-2 (complete or almost complete loss of uptake activity), Groups 3-6 (impaired or altered function) and Group 7 (no significant functional changes) (Supplementary Fig. 2a). Strikingly, variants in Groups 1-2 were observed exclusively in patients and not in controls (13 vs. 0), whereas the distributions for Groups 3-6 and Group 7 were 24 vs. 4 and 177 vs. 69, respectively (Supplementary Fig. 2b, c).

### Surface and total expression of DAT variants assessed by ELISA

To better understand the mechanism underlying the observed functional changes, we directly determined surface expression alongside total, cellular DAT expression of the variants by use of an enzyme-linked immunosorbent assay (ELISA) (Fig. 4 and Supplementary Table 1). Total and surface DAT expression were assessed on glyoxal-fixed cells using two primary antibodies: MAB369, targeting the DAT intracellular N-terminus, was used on permeabilized cells for total expression, and sc-32259, targeting EL2, was used on intact cells for surface expression (Fig. 4a-c)^40^.

**Figure 4.**
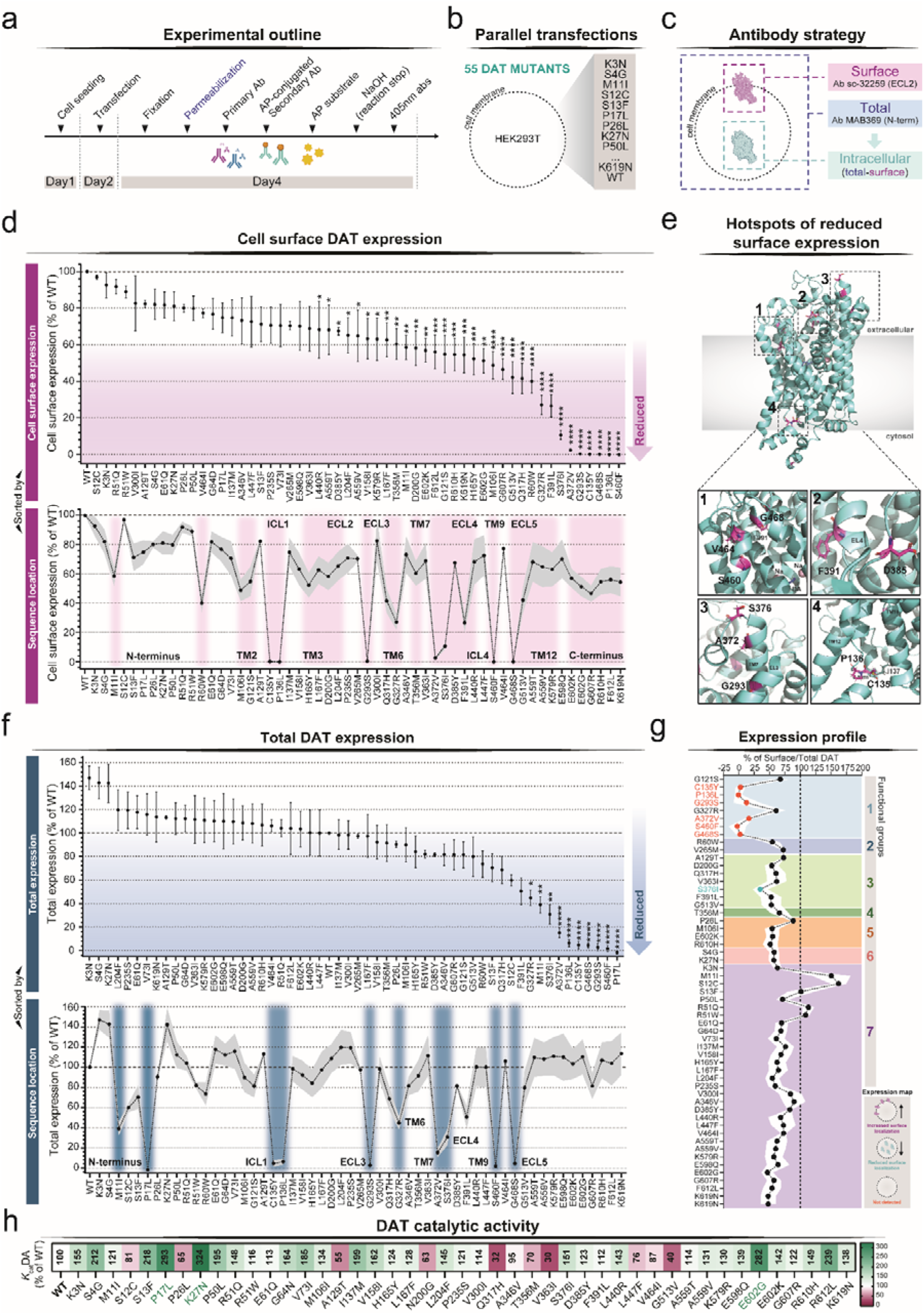
Cell-surface and whole-cell expression patterns of DAT mutants. **a** Schematic overview of the ELISA experimental outline. The permeabilization step (blue) denotes the additional step used to measure total DAT expression. **b** Graphical representation of the transfection setup. **c** Schematics of the antibody strategy adopted in the ELISA. **d** Cell-surface expression of DAT variants measured by ELISA, expressed as a percentage of WT surface expression. In the upper panel, DAT mutations are sorted according to expression levels. In the lower panel, the same data is mapped according to the position of each mutant in the DAT amino acid sequence. Pink areas refer to statistically significant values (one-way ANOVA with Holm-Šídák’ multiple comparisons test: *, p < 0.05; **, p < 0.01 ***, p < 0.001; ****, p < 0.0001). **e** Representation of the cryo-EM structure of the DA-bound human DAT (PDB: 8Y2D highlighting residues with strongly reduced cell-surface expression (magenta), as shown in panel **d** Numbered squares (1-4) magnify regions containing multiple mutations within this group. **f** Total expression of DAT mutants detected by ELISA, normalized to the total expression of the WT. The upper graph shows mutants sorted by expression levels, while the lower graph shows values according to their position in the DAT amino acid sequence. Light-blue areas indicate statistically significant results (one-way ANOVA with Holm-Šídák’ multiple comparisons test: *, p < 0.05; **, p < 0.01 ***, p < 0.001; ****, p < 0.0001). **g** Expression profile of DAT mutants calculated as surface-to-total ratio and sorted based on functional groups. Statistically significant mutants showing intracellular retention are displayed in turquoise, whereas mutants for which neither surface nor total expression was detected are indicated in orange. Statistical analysis: one-way ANOVA with Holm-Šídák’ multiple comparisons test. **h** Heat map of DAT catalytic activity (*K*_cat_DA) expressed as percentage of WT. Green mutants indicate statistically significant values (one-way ANOVA with Holm-Šídák’ multiple comparisons test: **, p < 0.01 ***, p < 0.001). Ab, antibody; AP, alkaline-phosphatase.

In the group of inactive variants (Group 1), we observed for C135Y, P136L, G293S, A372V, S460F and G468S loss of surface signal, suggesting that inactivity is caused by lack of cell surface expression (Fig. 4d, e and Supplementary Table 1). We also observed dramatically lower total expression for these six variants (Fig. 4f, g and Supplementary Table 1). For G327R, we observed markedly reduced surface expression (∼25% of WT) but only ∼50% reduction in total expression (Fig. 4d-g and Supplementary Table 1), consistent with a former study showing possible retention in the endoplasmic reticulum^41^. Similarly, G121S showed ∼50% reduction in surface expression and unchanged total expression compared to WT (Fig. 4d-g and Supplementary Table 1).

In Group 2, which was defined by loss of measurable DA uptake but partially retained DG3-80 binding (Table 2), R60W displayed reduced surface expression (∼40% of WT) with no difference in total expression, while V265M showed no significant reduction in surface and total expression. This adds further support that the loss of uptake is due to impaired catalytic activity combined with partial intracellular retention of R60W (Fig. 4d-g and Supplementary Table 1).

In Group 3, A129T and V363I displayed non-significant changes in surface expression, suggesting that the reduced uptake is mainly due to impaired catalytic activity (Fig. 4d-g and Supplementary Table 1). Q317H and G513V showed reduced surface expression (∼40% of WT for both variants) (Fig. 4d-g and Supplementary Table 1) but uptake capacity was even lower (Fig. 2 and Table 2), supporting both intracellular retention and impaired catalytic activity of this mutant. For D200G, S376I and F391L, the loss of surface expression (∼60%, ∼10% and ∼30% of WT, respectively; Fig. 4d-g, Supplementary Table 1) largely paralleled loss of uptake (∼36%, ∼16% and ∼30% of WT, respectively; Table 2), suggesting that intracellular retention is a dominating reason for the impaired uptake.

T356M (Group 4), characterized by decreased DA affinity (Table 2), demonstrated modest though significant reduction in surface expression with no change in total expression (Fig. 4d-g, Supplementary Table 1), adding further support to primarily a direct or indirect effect of the mutations on the substrate-transporter interaction.

Within Group 5, M106I, E602K and R610H showed ∼50% reductions in surface expression with no change in total expression (Fig. 4d-g, Supplementary Table 1), consistent with the lowered *B*_max_ for DG3-80 (Table 2), and pointing to intracellular retention as the cause of reduced activity. P26L, however, displayed no significant changes in surface and total expression (Supplementary Table 1).

The N-terminal variants in Group 6 (S4G, P17L and K27N) displayed surface and total ELISA signals >70% of WT. Together with increased uptake capacity, this resulted in significantly elevated catalytic activity for P17L and K27N (∼290% and ∼320% of WT, respectively; Fig. 4h, Supplementary Table 1) and a non-significant tendency towards higher *K*_cat_ for S4G (∼210% of WT; Supplementary Table 1), For P17L, the total expression signal was absent while surface expression was retained (∼75% of WT). A possible explanation is that this mutant is part of the epitope recognized by the antibody (MAB369, see below).

As expected for Group 7 variants, with no significant phenotype in the uptake and DG3-80 binding assays, a large fraction of mutants, including K3N, S12C, S13F, P50L, R51Q, R51W, E61Q, G64D, V73I, I137M, P235S, V300I, A346V, L447F, V464 and E598Q, showed no significant changes in surface or total expression (Fig. 4d-g, Supplementary Table 1). However, we detected a diminished surface signal for M11I, V158I, H165Y, L167F, L204F, D385Y, L440R, A559T, A559V, K579R, E602G, G607R, F612L and K619N (Fig. 4d, Supplementary Table 1). For E602G, the lowered surface expression (∼50% of WT) together with ∼145% uptake, resulted in significantly increased catalytic activity (*K*_cat_ ∼280% of WT; Fig. 4h, Supplementary Table 1), supporting a gain-of-function phenotype.

To establish that absence of signal in the ELISA, together with no detectable uptake and DG3-80 binding, would not arise from reduced quality of plasmids or transfection efficiency, we performed western blots on C135Y, P136L, G293S, A372V, S460F and G468S, utilizing MAB369 primary antibody on cell lysates from transiently transfected cells. We observed for all these mutations clear bands at ∼55 kDa, i.e. the expected molecular weight of non-glycosylated hDAT monomers, while none of them had a band corresponding to the mature, glycosylated form of DAT (i.e. close to 75 kDa), further substantiating the hypothesis of folding and/or trafficking deficits in these variants (Supplementary Fig. 3a)^42^. Of note, we also tested P17L. However, as this mutant did not seem to bind MAB369 in the ELISA studies, despite being otherwise kinetically functional, we used the antibody targeting ECL2 instead (sc-32259). Indeed, both 55 kDa and 75 kDa bands were detected, confirming expression of P17L (Supplementary Fig. 3b).

### Confocal imaging of selected mutants

To further investigate whether alterations in uptake and cocaine-analogue binding (Fig. 2, Fig. 3 and Table 2) stemmed from impaired DAT trafficking/delivery to the plasma membrane, we performed confocal imaging on variants displaying highly impaired DA uptake (R60W, G121S, C135Y, P136L, V265M, G293S, Q317H, G327R, A372V, S376I, S460F and G468S). We carried out separate, double staining of DAT using the sc-32259 antibody targeting EL2 to visualize plasma membrane expression and, upon permeabilization, the MAB369 antibody, targeting the DAT N-terminus to visualize total expression of DAT. Subsequent image processing revealed two phenotypes. One subgroup (G121S, Q317H and S376I) exhibited clear surface stain, along with positive MAB369 signal, supporting that the impaired uptake and DG3-80 binding for these mutants do not merely arise from reduction in cell surface expression (Supplementary Fig. 4). For R60W, P136L, V265M and G327R, we observed diminished, yet detectable, cell surface staining (Supplementary Fig. 4). Conversely, surface signal was completely abolished for C135Y, G293S, A372V, S460F and G468S; however, an intracellular signal was still clearly visible consistent with intracellular retention of these variants (Supplementary Fig. 4).

### Assessment of amphetamine-mediated DA efflux of DAT coding variants

DAT is a primary target for amphetamine (AMPH), which is both an illicit psychostimulant and a prescribed treatment for ADHD^43,44^. As a DAT substrate, AMPH competes with DA uptake and promotes an efflux-prone transporter state that includes DAT phosphorylation^45–47^. To assess AMPH-induced DA efflux, we used GRAB_DA2M_ “sniffer cells”, HEK293 cells expressing a genetically encoded DA sensor (Fig. 5a)^48^. Cells were transfected with DAT WT, DAT mutants, or empty vector (Fig. 5b). Following loading with 1 μM DA for 15 min, the fluorescent signal (indicative of extracellular DA) was ∼30% lower in WT-transfected cells than in mock-transfected cells, as a likely result of DAT-mediated uptake of the added DA into the cells (Fig. 5d)^48^. In line with this, we observed fluorescence changes that varied according to the uptake activity of each mutant (selected mutants included in Fig. 5d). After two washes and a 20-min incubation, cells were challenged with 30 μM D-AMPH and fluorescence monitored for an additional 15 min (Fig. 5b,e). WT DAT produced a ∼40% increase in extracellular DA, whereas mutant responses varied considerably (Fig. 5c,e).

**Figure 5.**
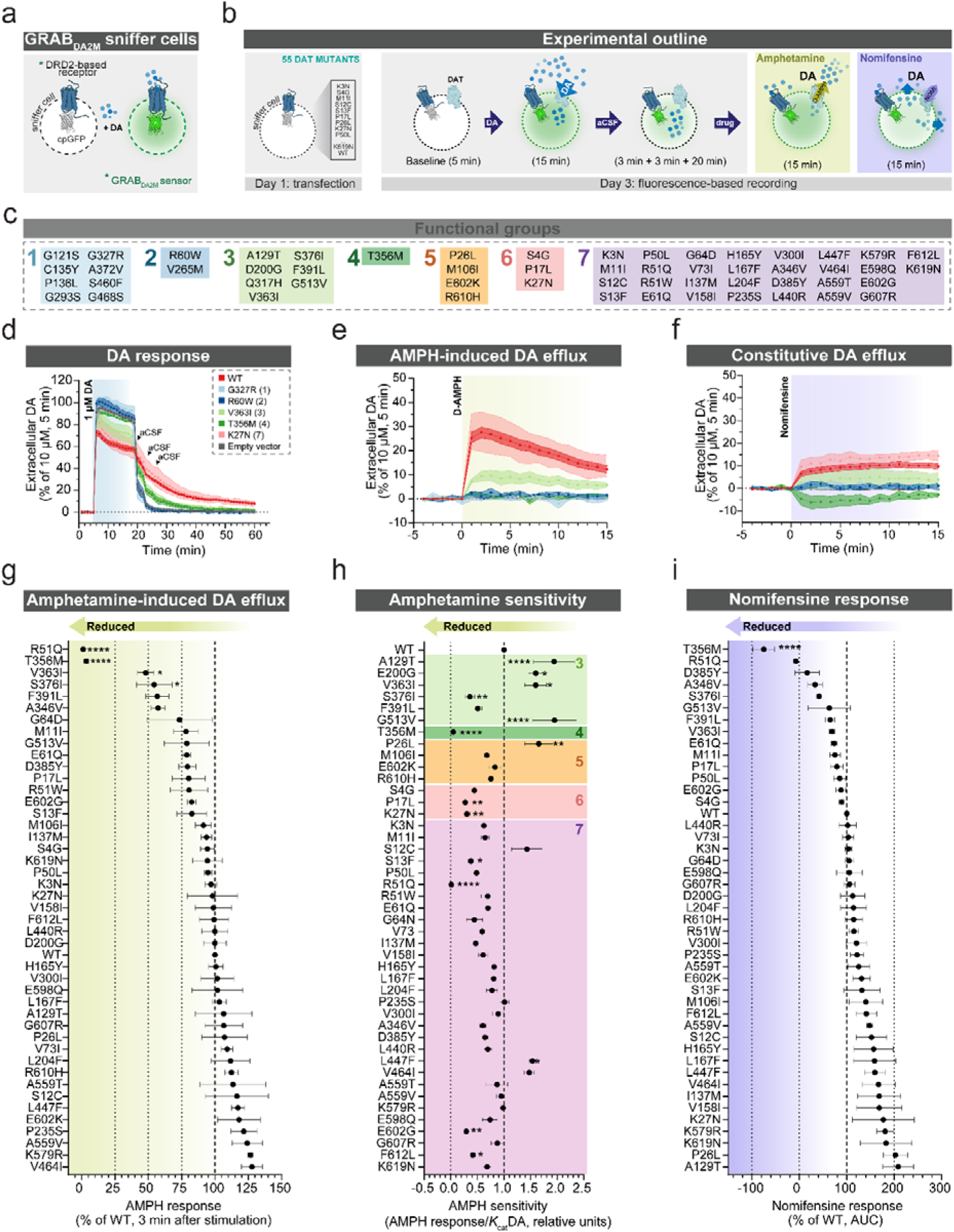
Evaluation of D-AMPH-elicited DA efflux and constitutive efflux through GRAB_DA2M_ sniffer cells assay. **a** Schematic illustration of GRAB_DA2M_ sniffer cells assay. **b** Experimental workflow (see “Materials and methods” for detailed procedures). **c** List of DAT mutants grouped and color-coded according to their functional groups. **d** DA fluctuations recorded using GRAB_DA2M_ sniffer cells, transiently and separately transfected with the WT, pcDNA3.1 (empty vector), K27N, R60W, G327R, T356M and V363I, here selected as representative mutants. Values are normalized to the signal measured 5 minutes after addition of 10 µM DA. **e** AMPH response of WT and selected mutants measured on GRAB_DA2M_ sniffer cells, pre-loaded with 1 µM DA. The assay is performed as shown in panel **b**, with addition of 30 µM D-AMPH at minute 47. Data are normalized to 5 minutes after 10 µM DA administration and result from at least three independent experiments, each performed in duplicates. **f** Measurement of DAT-mediated constitutive DA efflux of WT and representative mutants on GRAB_DA2M_ sniffer cells, pre-loaded with 10 µM DA. The assay is performed as shown in panel **b**, with addition of 10 µM nomifensine at minute 47. Traces begin 5 minutes prior to addition of nomifensine and are normalized to the signal measured 5 minutes after 10 µM DA administration. Data represent at least three independent experiments, each performed in duplicates. **g** AMPH-induced DA efflux of DAT mutants measured on GRAB_DA2M_ sniffer cells 3 min after AMPH addition. Data are represented as mean ± S.E.M. of 3 to 5 independent experiments, each performed in duplicates and normalized to the WT response. **h** AMPH sensitivity of DAT mutants (AMPH response/*K*_cat_DA), sorted by functional group. **i** Constitutive DA efflux measured in response to 10 µM nomifensine. Values represent the area under the curve (AUC) of 3 to 5 independent experiments, each performed in duplicates and are normalized to the WT response. Statistical test in **g**-**i**: one-way ANOVA with Holm-Šídák’ multiple comparisons test.

Because AMPH-induced efflux in this assay depends on prior DAT-mediated DA loading, all variants lacking detectable DA uptake were excluded (R60W, G121S, C135Y, P136L, V265M, G293S, G327R, A372V, S460F, and G468S; Groups 1–2). Q317H (Group 3) was also excluded because uptake was only ∼13% of WT (Table 2).

Among the remaining variants, AMPH responses differed substantially (Fig. 5g). Peak responses were reduced by ∼50% for V363I and ∼45% for S376I (Group 3; Fig. 5g; Supplementary Table 2). Interestingly, AMPH elicited no response in R51Q (Group 7), despite its otherwise functional profile, or in T356M (group 4; Fig. 5g; Supplementary Table 2). For T356M, this likely reflects constitutive DA efflux that limits DAT-dependent DA uptake^16,49^.

Normalization of AMPH responses to DA transport capacity (AMPH response/*K*_cat_DA) identified significantly increased AMPH sensitivity for A129T, D200G, V363I and G513V (Group 3), P26L (Group 5), and L447F (Group 7) (Fig. 5h and Supplementary Table 2). Conversely, all gain-of-function variants (P17L and K27N in Group 6 and E602G in Group 7), as well as S376I (Group 3), S13F, and F612L (Group 7), showed reduced sensitivity to AMPH (Fig. 5h and Supplementary Table 2).

### Determination of DAT-mediated constitutive DA efflux

Few DAT mutations have been reported to promote an efflux-prone state, resulting in anomalous or constitutive dopamine (DA) efflux (ADE). This includes the disease-associated variants A559V, T356M, and D421N^13,14,16,18^. These variants exhibit diverse functional profiles ranging from near-normal uptake activity to severe impairment^14–16,19^. We reasoned that our large cohort of disease variants could provide an opportunity to assess the prevalence of this atypical molecular phenotype.

To this end, we again used GRAB_DA2M_ sniffer cells, previously shown to detect ADE in T356M^48^. Cells were loaded with 10 µM DA, washed, and challenged with the DAT inhibitor nomifensine (Fig. 5b, f). Previously, we reported that blockade of WT induced a slow increase in extracellular DA, consistent with DAT-independent leakage of DA from pre-loaded cells and blocked reuptake of the leaked DA. In contrast, T356M produced a decrease upon nomifensine application, consistent with blockade of mutant-mediated ADE (Fig. 5f)^48^. Additional support for an efflux phenotype was the elevated extracellular DA in the GRAB_DA2M_ assay during aCSF wash steps, relative to uptake-inactive mutants (Fig. 5d). Representative GRAB_DA2M_ traces following nomifensine application are shown in Fig. 5f.

All other tested variants showed non-significant changes in response to nomifensine (Fig. 5i). Notably, A559V, previously suggested to mediate constitutive efflux, displayed a larger, rather than reduced, nomifensine response (∼150% of WT, not significant), while A559T exhibited a response of ∼125% of WT, arguing against detectable ADE under the assay conditions (Fig. 5i; Supplementary Table 2)^18,50^.

To exclude that limited assay sensitivity obscured potential efflux phenotypes, we also employed a classical superfusion efflux assay based on pre-loading the cells using [^3^H]MPP^+^, a membrane impermeable DAT substrate^45^. A346V, V363I and D385Y (plus T356M as control) were tested because, compared to WT and like T356M, they showed varying degrees of residual uptake capacity, impaired AMPH-response, reduced response to nomifensine and/or increased p*K*_m_DA. Nomifensine did not significantly alter [^3^H]MPP^+^ release for any variant, although T356M showed a trend toward a reduced response (Supplementary Fig. 5a, b). However, basal extracellular [^3^H]MPP^+^ levels were significantly elevated for A346V and T356M (Supplementary Fig. 5c-e), suggesting that A346V, like T356M, may support low-level constitutive efflux. We also tested AMPH showing, in agreement with the sniffer cell assay, significantly reduced AMPH-induced [^3^H]MPP^+^ efflux for A346V, V363I and D385Y and, as expected based on previous findings, reduced AMPH-induced [^3^H]MPP^+^ efflux for T356M (Supplementary Fig. 5f-h)^16,49^.

### Molecular dynamics simulation on selected mutants

To investigate the structural basis of the observed molecular phenotypes, we performed molecular dynamics (MD) simulations on four selected mutants, S460F (Group 1) and Q317H, V363I and S376I (Group 3), each in both (i) DA-free and (ii) DA-bound states. We assessed the effects of the mutations on the stability of the outward-open conformation, substrate-induced occlusion and inner gate stability (Fig. 6a-h). Degree of occlusion was quantified by measuring the distance between the upper portion of TM9 (scaffold domain) and TM6a (bundle domain) (Fig. 6a). In the presence of DA, all mutants showed impaired occlusion, with S460F, Q317H and V363I remaining in an outward-facing conformation, and S376I adopting a conformation closer to the occluded state (Fig. 6a, b, f).

**Figure 6.**
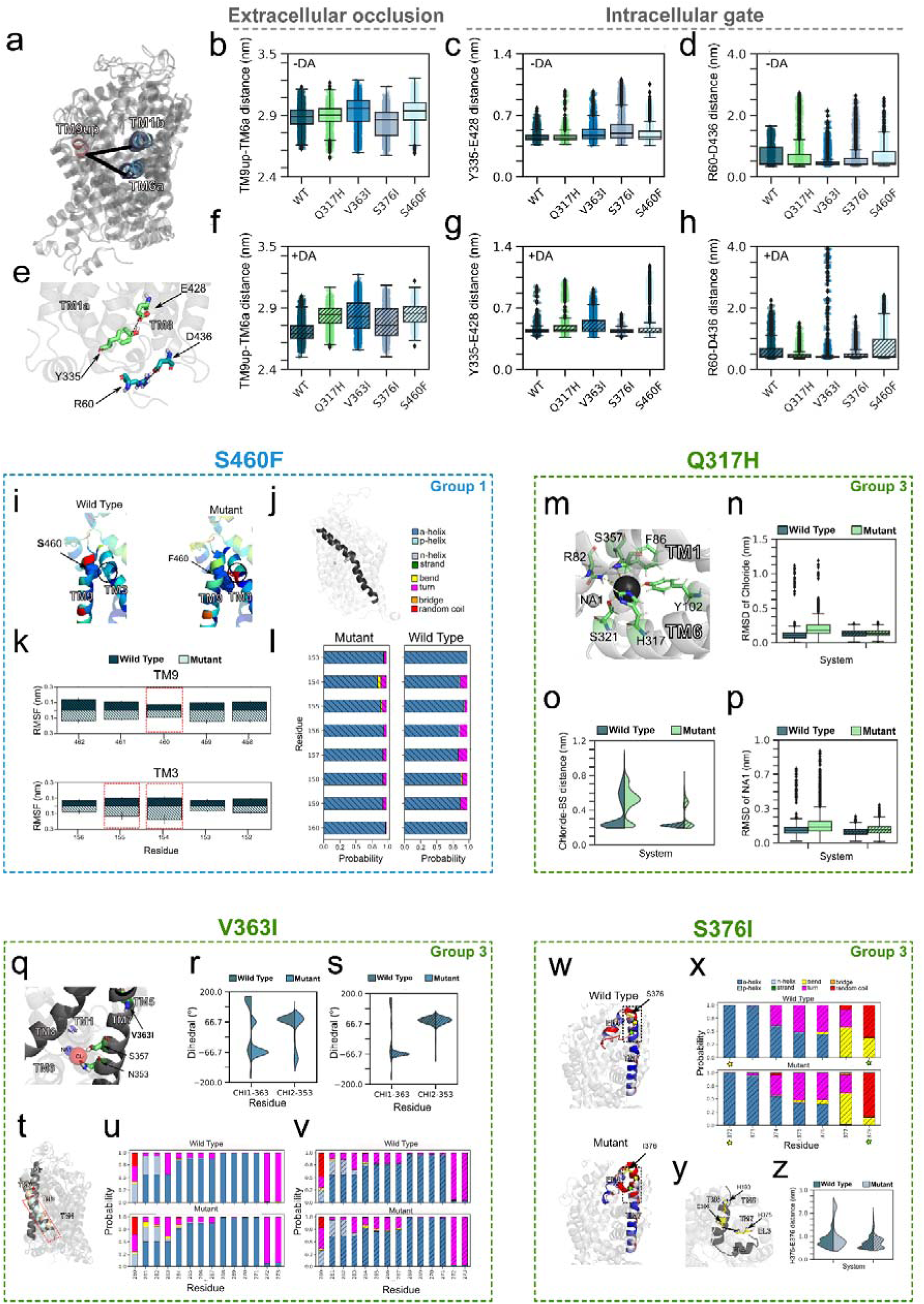
Global and local structural changes of selected DAT variants. **a**, **e** Structural measurements used to assess extracellular occlusion (**a**) and intracellular gating (**e**). **a** Superposition of representative DA-free and DA-bound frames, illustrating TM9up relative to TM6a and TM1b. **e** Close-up on the intracellular gate formed by Y335-E428 and R60-D436 interactions. **b**-**d**, **f**-**h** Boxplots displaying distribution of TM9up-TM6a (**b, f**), Y335-E428 (**c**, **g**) and R60-D436 (**d**, **h**) from all replicates for WT, Q317H, V363I, S376I, and S460F, under DA-free and DA-bound states. Boxes indicate the interquartile range (IQR), the center line represents the median, whiskers extend 1.5 up the IQR and grey squares indicate outliers. **S460F: i** RMSF of each residue in TM9 and TM3 projected onto the DA-bound system for both WT and mutant, colored from lower (blue) to higher (red) values. **j** TM3 location in the structure. **k** Average RMSF of TM9 and TM3 residues across replicates. Red dashed lines highlight the mutated residue and neighboring TM3 residues. **l** Average TM3 secondary structure changes in the DA-bound WT and mutant. **Q317H: m** Close-up view of Na1 and chloride binding sites close to Q317. **n**, **p** Boxplots of chloride (**n**) and Na1 (**p**) RMSD in the DA-bound and DA-free states. **o** Average distance distribution of chloride-coordinating residues. **V363I: q** Local environment surrounding V363. **r**, **s** Dihedral angle (chi1) distributions for residues 363 and 353 under DA-free (**r**) and DA-bound (**s**) conditions. **t** Superposition of WT (pink) and mutant (light blue) DA-free structures, highlighting TM5. **u**, **v** Stacked bar charts of average secondary structure changes WT (top) and mutant (bottom) under DA-free (**u**) and DA-bound (**v**) states. **S376I: w** RMSF mapped onto TM7 and EL4 in the DA-bound system for both WT and mutant. **x** TM7 and EL4 secondary structure changes (region highlighted with stars in **w**) in the DA-bound state. **y** Close up view of the zinc-binding site. **z** H375-E396 distance distribution of WT and mutant in presence and absence of DA.

To assess inner gate stability, we evaluated the stability of the salt bridge in the closed inner gate (R60-D436) and the distance between Y335 and E428 in the core of the inner vestibule (Fig. 6e). In the WT, the Y335-E428 distance was consistent with a closed inner gate in both the DA-free and DA-bound states. By contrast, all mutations showed perturbed hydrogen bond and/or salt bridge distance distributions. For both V363I and S376I, there was in the DA-free state a trend towards reduced Y335-E428 hydrogen bond stability together with increased R60-D436 salt bridge solidity (Fig. 6c, d), while Q317H also strengthened the salt bridge, especially in the DA-bound state (Fig. 6c, d). In the DA-bound state, V363I was the only mutant destabilizing the Y335-E428 hydrogen bond, whereas Q317H, V363I and S376I all stabilized the R60-D436 salt bridge relative to WT. Summarized, the mutations induced global conformational changes, predominantly trapping DAT in an outward-open state with accompanying destabilization of the inner gate.

To further delve into the structural effects caused by the four mutants, we examined local perturbations surrounding each variant. S460F is located in the extracellular region of TM9 adjacent to TM3 (Fig. 6i), leading us to hypothesize that S460F could affect the neighboring TM3. Root mean squared fluctuation (RMSF) analysis revealed increased flexibility in TM3 residues 154-155 facing the S460F side chain (Fig. 6k, l), accompanied by stiffening of the TM3 structural kink (Fig. 6i, j). These structural changes may impair transporter folding/surface targeting, consistent with the absence of uptake activity and detectable surface expression (Table 2 and Supplementary Table 1). Supporting this interpretation, western blotting detected S460F predominantly in its immature, non-glycosylated form (Supplementary Fig. 3). Moreover, the simulation indicated that, if adequately folded and expressed, the mutation would still perturb transporter conformational dynamics, favoring an outward-facing state.

Q317 directly engages in chloride coordination via its amide group and when replaced by histidine in the Q317H mutant, the interaction with chloride is maintained, but via the histidine’s amine group (Fig. 6m). MD simulations revealed a displacement of chloride from its binding site in both DA-free and DA-bound states compared to WT (Fig. 6n, o). Given the proximity of chloride to the Na1 sodium site, we assessed whether instability of chloride affected sodium coordination. Indeed, Na1 deviated from its initial position in both conditions, with the effect being more pronounced in the DA-free state (Fig. 6p). Thus, Q317H likely impairs transport by perturbing both chloride and Na1 sodium binding.

V363I is located in the outer part of TM7 at the TM5 interface, two turns from S357 and N353 (Fig. 6q), which contribute to the sodium (Na1) and chloride binding site^1,24^. MD simulation showed that the extra methyl group introduced by isoleucine reduces V363 rotation flexibility under both DA-free and DA-bound conditions (Fig. 6r, s). The loss of entropy associated with the reduction of residue mobility was compensated by shifting the dihedral angle of N353 (Fig. 6r, s). When the chi1 dihedral angle of N353 assumes values equal to +66.7°, its amide group faces chloride while its carbonyl oxygen faces Na1. The inversion of this dihedral leads to strong repulsive forces, which affects sodium and chloride binding. Additionally, reduced entropy increases mobility in the lower TM5 region, promoting destabilization of secondary structure around residues 260-264 (Fig. 6t-v). In summary, V363I indirectly affects sodium and chloride coordination through allosteric effects, distinct from the direct, orthosteric perturbation observed for Q317H.

S376I is located at the outer end of TM7 at the transition to EL4 and close to the zinc binding site (E396, H375, and H193)^51,52^. Replacement of the polar serine with hydrophobic isoleucine increased mobility of the outer TM7 region (Fig. 6w), accompanied by a shift from a structured (helix) to a less structured (turn or bend) conformations of both TM7 and EL4 (Fig. 6x). Interestingly, these changes influenced the zinc binding site organization (Fig. 6y, z), resulting in a multimodal distribution of distances between the side chains of H375 and E396 or H193, compared to WT (Fig. 6y, z). Thus, S376I destabilizes the outer TM7 region, which conceivably may contribute to reduced transporter surface expression and the associated reductions in uptake and DG3-80 binding.

## DISCUSSION

Given the central role of DAT in balancing vesicular, cytoplasmic and extraneuronal DA, DAT dysfunction is highly relevant in the context of neuropsychiatric disorders where DAergic dysregulation is suspected to be involved. Amongst such conditions are the ones included in our case-cohort sample, encompassing patients diagnosed with at least one of these disorders: ADHD, ASD, BD, SCZ or depression^22^. Indeed, the importance of loss-of-function intolerance in *SLC6A3* has been strongly supported by the identification of disease-associated ultra-rare or rare variants in the coding regions that associate with a spectrum of diseased states. These range from infantile parkinsonism dystonia (DTDS) to neuropsychiatric disease, conceivably dependent on the molecular phenotype of the mutants, including dominant negative properties, and whether the patients are homozygous or heterozygous for the genetic variant^8–12,53^.

Here, we carried out a systematic and comprehensive analysis of multiple DAT variants by exploiting a cohort of Danish psychiatric patients^22^. Of the selected 55 DAT missense mutants, 24 variants have been reported previously as disease-associated in the literature or in the ClinVar database, leaving 31 variants as new (Table 1). Notably, when interrogating PolyPhen-2 to predict the structural damage imposed by a specific mutation, 31 of the 55 mutants scored as either “possibly damaging” or “damaging”, while 24 were considered “benign”. Although we did observe overlap, e.g. the variants in Group 1 all scored as damaging by PolyPhen-2 consistent with complete loss of function in our assays, there were also clear discrepancies with e.g. D200G, F391L and S376I (Group 3) being predicted to be benign yet showing major loss of function in our assays. Furthermore, PolyPhen-2 is unlikely to predict possibly disease-promoting, gain-of-function phenotypes such as constitutive efflux^54^. These findings underscore the importance of experimentally validating *in silico* predictions of variant pathogenicity and thereby directly addressing a fundamental challenge in human neuroscience: how to translate the rapidly growing catalog of rare neuropsychiatric variants into mechanistic understanding.

Functional profiling classified the variants into seven distinct phenotypic groups according to their uptake and binding characteristics. Intriguingly, the Group 1 and 2 variants, which showed either complete or marked loss of uptake function, were found exclusively among cases, further supporting a link between transporter dysfunction and disease. Moreover, the combined assessment of uptake activity, inhibitor binding, surface expression by ELISA, and cellular distribution by confocal imaging enabled us to identify convergent mechanisms responsible for the functional impairment of the Group 1-2 variants. For the Group 1 variants C135Y, P136L, G293S, A372V, S460F and G468S, the loss of uptake was accompanied by a similar loss of surface expression according both to the ELISA assay and confocal imaging supporting a major trafficking defect possibly involving intracellular retention or increased endocytosis of the mutant transporters. MD simulations of S460F supported this notion by showing how insertion of a phenylalanine side chain in position 460 caused major secondary structure perturbations in TM3, likely impairing transporter folding/surface targeting.

In contrast, the loss of function for G121S and G327R was likely related to both a direct impairment of catalytic activity and a trafficking defect, because their residual surface expression was higher than their uptake capacity. Similarly, for R60W and V265M in Group 2, the functional impairment was conceivably mainly caused by defective catalytic activity. A similar scenario was seen for the variants A129T, Q317H and V363I in Group 3. Collectively, the observed functional impairments could be attributed to defined molecular mechanisms involving disrupted substrate and ion coordination or impaired conformational transitions during the transport cycle. The R60W variant, for example, is intriguing, as introduction of a tryptophan at this site is predicted to disrupt the well-characterized interaction network that constitutes the intracellular gate of the transporter^32,55^. Q317H is also of specific interest because the glutamine at this position is a critical component of the chloride binding site. Indeed, simulations supported perturbation of not only chloride binding but secondarily also of the Na1 in this mutant, explaining the complete loss of function observed. For V363I, MD simulations pointed to an indirect allosteric perturbation of the sodium and chloride binding sites caused by the single additional methyl group in the mutant side chain as a conceivable explanation for the decreased uptake.

For the remaining mutants in Group 3 (D200G, S376I F391L and G513V), lowered *V*_max_ values most likely reflected a trafficking defect although less severe compared to that seen for the intracellular retained mutants in Group 1. MD simulations on S376I supported that introduction of an isoleucine in position 376 on the outer face of the transporter adjacent to the zinc-coordinating residue H375 in DAT not only perturbed the zinc binding site itself but also destabilized the outer part of TM7 possibly leading to folding impairment and possibly therefore reduced surface expression^51,52,56^. Equivalent indirect mechanisms can be envisioned for D200G, F391L and G513V. G513V, for example, is localized in the fourth intracellular loop, where replacement of the highly conserved glycine with valine would be expected to perturb secondary structure that in turn could have a destabilizing effect interfering with folding and/or surface targeting,

T356M (Group 4) is a particularly noteworthy variant, as we confirmed the previously reported constitutive DA efflux phenotype^14,16^. However, the structural basis underlying this efflux remains poorly understood. Our comprehensive analyses suggest that this phenotype is rare, as none of the other variants exhibited T356M-like phenotype in our sniffer-cell assay. However, using the more sensitive [³H]MPP⁺ superfusion assay, we detected a significant increase in basal efflux for A346V, suggesting that this variant may also mediate constitutive efflux, albeit to a markedly lesser degree than T356M. We cannot exclude that additional variants display similarly modest (as A346V) or mechanistically distinct efflux phenotypes, which may have gone undetected. For example, we were unable to reproduce the proposed efflux phenotype of the variant A559V^18^. This discrepancy may reflect the reported evidence suggesting that A559V-mediated DA efflux depends on dopamine D2 receptor activation and post-translational modifications including phosphorylation^57^. Indeed, receptor expression and/or signaling pathways may differ across heterologous expression systems, therefore influencing the detection of the phenotype^29,50,57^. Further studies outside the scope of this work will be required to resolve these issues.

Our analysis did not only identify loss-of-function mutations but also putative gain-of-function variants. This includes residues in Group 6 located on the DAT N-terminus and C-terminus. P17L and K27N in the N-terminus showed significantly increased maximal transport velocity relative to surface expression, while S4G showed a trend toward enhanced transport capacity. Furthermore, the C-terminal variant E602G in Group 7 exhibited increased catalytic capacity. The pathophysiological consequences of enhanced versus reduced transport capacity remain uncertain yet it is interesting to note that gain-of-function variants have also been identified in the homologous serotonin transporter (SERT), including the ASD- and OCD-associated N-terminal variant G56A^58^. Interpreting the putative mechanistic implications of N-terminal mutations, as well as other cytosolic domains, requires consideration of how the variants might alter post-translational regulation, including phosphorylation, palmitoylation and ubiquitination (K27 is a major ubiquitination site), as well as interactions with DAT-associated proteins^39,59–61^. In turn such changes may not only alter transporter trafficking, but also directly affect its function^62–65^.

For the variants in Group 7, we observed overall more modest effects across the functional assays performed. While some of these variants, despite being found in patients, may have absent to only marginal effect on DAT function and consequently contribute only partially to the observed disease phenotypes, it is also possible that crucial disease-relevant mechanisms are not fully captured by screening the mutants in an overexpression system. An illustrative example is K619N, located in the critical C-terminal PDZ (PSD-95/Discs-large/ZO-1 homology)–binding motif of DAT and associated with a spectrum of disease-phenotypes including psychiatric disease and atypical DTDS^12^. In HEK293 cells, both in the present study and in our previous work, K619N displayed relatively modest impairment^14^. However, more detailed investigation revealed a dominant-negative effect on the WT transporter counterpart in relation to surface targeting, and *in vivo* expression models in *Drosophila* and mice demonstrated direct effects on dopamine-directed behaviors, as well as dominant-negative interference with WT transporter function^12^. Such detailed investigations of individual variants lie beyond the scope of this study; however, the findings highlight the importance of cellular contexts, expression systems and regulatory mechanisms when evaluating the disease-impact of clinical DAT variants.

In summary, this study provides a comprehensive framework for understanding how rare and ultra-rare patient-derived DAT missense variants disrupt transporter function. By integrating functional, cellular and structural analyses, we show that these variants converge on a limited number of molecular mechanisms, including defective trafficking, impaired catalytic activity and altered transporter dynamics. Our findings establish thereby a valuable resource for interpreting the functional consequences of rare SLC6A3 variants identified in psychiatric patients and provide new insight into how DAT dysfunction may contribute to the pathophysiology of major neuropsychiatric disorders.

## METHODS

### Source of genetic and diagnostic information

Exome sequencing data at the basis of our investigation was obtained from the Lundbeck Foundation Initiative for Integrated Psychiatric Research (iPSYCH) which carried out an extensive study based on the Danish population born between 1^st^ of May 1981 and 31^st^ of December 2005, including a starting sample of 1,472,762 people, as described elsewhere^22^. Our investigation is based on a case-control sample nested within the iPSYCH2012 case-cohort study, whereby individuals diagnosed with at least one of five major mental disorders, i.e. ADHD, affective disorder, ASD, BD and SCZ, were classified as cases (N = 57,377), according to the International Classification of Disease, 10^th^ revision. In this cohort, 30,000 people were randomly assigned to the control group. Using blood samples taken 4-7 days after birth and retrieved form the Danish Neonatal Screening Biobank, a portion of the original group, featuring 19,851 individuals and comprising 5,554 controls, underwent array genotyping and variant annotation. Disease groups are represented in the cohort as follows: ADHD: N=5040, affective disorder: N=2967, ASD: N=5694, SCZ: N=3686, and BD: N=1528. Quality control assessment was applied to the original sample, with 77,639 out of 86,198 individuals delivered to the next steps. This study is rooted in the latest QC-refined iPSYCH2012 dataset, where 83 missense mutations in the *SLC6A3* gene have been identified across the study group. For our investigation, we selected a sub-group of missense DAT variants with preferential segregation to the comprehensive disease category (patients/controls >1). The Danish Data Protection Agency, the Danish Scientific Ethics Committee, the Danish Health Authority and the Danish Neonatal Screening Biobank Steering Committee approved the iPSYCH study.

### Cell culture and transfection

DNA constructs used for transfections were purchased from BioCat GmbH. Each point mutation was introduced on a pcDNA3.1 vector carrying human DAT WT cDNA, and subsequently verified by sequencing. [^3^H]DA uptake, DG3-80 binding, ELISA and confocal images acquisition were performed on transiently transfected human embryonic kidney (HEK293) cells, maintained in Dulbecco’s modified Eagle’s medium (DMEM), supplemented with 10% (v/v) fetal bovine serum (FBS, Thermo Fisher Scientific) and 1% (v/v) penicillin/streptomycin. Sniffer cells were cultured in DMEM supplemented with 10% (v/v) FBS, 1% (v/v) penicillin/streptomycin, 200 µg/mL Hygromycin B (Sigma) and 15 µg/mL Blasticidin (Gibco), as previously described^48^. All cells were maintained in cell culture-treated flasks, at 37°C and 5% CO_2_-supplied incubators. Two days before performing the assay, cells were transfected with DAT WT or DAT mutants, using Lipofectamine2000 (Invitrogen) and 3 µL/µg DNA, then seeded onto poly-L-ornithine-coated plates. Cell cultures were routinely tested using the MycoAlert detection kit (catalog #LT07-418, Lonza) assay to ensure that they were free from mycoplasma contamination.

For stable cell-line generation (used for the superfusion release assay), HEK293 cells were transfected using a reverse transfection protocol. In a tube, 50 µL serum-free Opti-MEM (Thermo Fisher Scientific) was mixed with Lipofectamine 2000 transfection reagent according to the manufacturer’s protocol. In a separate tube, 50 µL serum-free Opti-MEM was mixed with 2 µg of plasmid DNA. Transfection mix was prepared by mixing both tubes and incubation at room temperature for 5 min. Cells were seeded into a 6-well plate with 500 µL/well. Cells were subsequently transfected by adding 100 µL transfection mixture for 6 hours, then changing the medium to DMEM supplemented with 10% FBS. Selection was carried out over two weeks, using DMEM supplemented with 10% FBS and 150 µg/mL Geneticin and tested for expression via qPCR. After selection, cells were maintained in DMEM supplemented with 10% FBS and 50 µg/mL Geneticin at 37°C in humidified atmosphere with 5% CO_2_.

### [^3^H]DA uptake

[^3^H]DA saturation uptake experiments were conducted in whole-cell setup, in 96-well plates (catalog #3610, Corning Life Sciences) as described previously^14^. Briefly, two days after transfection, HEK293 cells were washed once in uptake buffer (120 mM NaCl, 5 mM KCl, 1.2 mM CaCl_2_, 1.2 mM MgSO_4_ and 25 mM HEPES, supplemented with 5 mM D-glucose, 1 mM ascorbic acid; pH 7.40) and equilibrated in uptake buffer for 10 min at room temperature. Saturation uptake was carried out using serial 1:2 dilutions (from 6.4 µM to 0.05 µM) of [2,5,6-^3^H]DA (PerkinElmer Life Sciences) and unlabeled DA (Sigma-Aldrich), kept in constant ratio. After 5min, uptake was stopped by washing cells twice in ice-cold buffer. After addition of scintillation fluid to each well, cells were left on mild shaking for 1h. Radioactivity was measured as counts per minute on a 2450 MicroBeta^2^ microplate counter (PerkinElmer, Waltham, MA, USA). Non-specific binding was determined using mock-transfected cells. Data are from a minimum of 3 independent replicates.

### DG3-80 saturation binding to DAT

Surface expression/binding features of DAT coding variants were tested using DG3-80, a fluorescent ligand that blocks DAT with low nanomolar affinity, as previously described^23^. Transiently transfected HEK293 cells expressing DAT WT or DAT mutants, were seeded on 96-well plates (catalog #6005689, PerkinElmer Life Science) at a density of ∼3×10^4^ cells/well. After 48 hours, cells were equilibrated in artificial cerebrospinal fluid (aCSF; 120 mM NaCl, 5 mM KCl, 2 mM CaCl_2_, 0.3 mM MgCl_2_ and 25 mM HEPES, supplemented with 5 mM D-glucose, 5.5 mM ascorbic acid and 0.2% BSA; pH 7.40) for 10 min at RT. Serial dilutions of DG3-80, ranging from 0.78 nM to 400 nM, were added to cells and left on incubation for 30 min. Binding was stopped by washing three times with aCSF. Fluorescence (540/580) was measured thereafter, using a POLARstar OMEGA plate reader (BMG Labtech), at 37°C. Nonspecific binding was determined on mock-transfected cells. Results are from at least 3 independent experiments.

### Measurement of total and cell surface expression of DAT by ELISA

DAT cell surface and total expression were quantified as reported previously^40^. In detail, 9.5×10^4^ HEK293 cells were seeded in 24-well plates previously coated with 0.02 mg/ml poly-L-ornithine hydrobromide (catalog #P3655, Sigma-Aldrich). After 24h, cells were transiently transfected with the indicated constructs using Lipofectamine2000 and incubated at 37°C for additional 48h. On the day of the experiment, cells were fixed for 30 min at 4°C using a cold Tris-buffered saline solution (TBS: 20 mM Tris, 150 mM NaCl, pH 8.0) containing 9% glyoxal, 8% acetic acid, and adjusted to a final pH of 4.0. Measurement of dopamine transporter cell surface expression was then carried out as previously described with some minor modifications^66^. Briefly, following two washes with TBS, non-specific binding was blocked by incubation with TBS containing 1% bovine serum albumin (BSA) for 30 min at room temperature. A rat monoclonal antibody raised against the second extracellular loop of DAT (catalog #sc-32259, Santa Cruz Biotechnology; 1:1000 in TBS-BSA 1%) was added and incubated for 60 min with gentle shaking at room temperature. Cells were then washed twice with TBS, blocked again with TBS-BSA 1% for 15 min and incubated with an alkaline phosphatase-conjugated rabbit polyclonal anti-rat IgG antibody (catalog #A6066, Sigma-Aldrich; dilution 1:10,000) for 60 min at room temperature with gentle shaking. For total expression of DAT, a rat monoclonal antibody raised against the intracellular *N*-terminal of DAT (catalog #MAB369, Sigma-Aldrich; dilution 1:750) was used as the primary antibody, and 0.02% saponin was added to the TBS buffer in all washing, blocking and antibody solutions in order to permeabilize the cells. After two more washes with TBS, the colorimetric reaction was initiated by the addition of 250 μL of a solution containing the alkaline phosphatase substrate (catalog #S0942, Sigma-Aldrich; 1 tablet/5 mL of solution) diluted in a diethanolamine solution (9.5% diethanolamine, 0.5 mM MgCl_2_, pH 9.8). The plates were incubated at 37°C until a yellow color appeared, and the reaction was stopped by the addition of 250 μL of NaOH 0.4M. A volume of 200 μL of the colorimetric reaction was then transferred into a 96-well plate and the absorbance at 405 nm was measured using the POLARstar Omega microplate reader. For each independent experiment, all conditions were carried out in triplicate (cell surface expression) or duplicate (total expression). The results are from 5 independent experiments.

### Western blotting

Expression patterns of selected DAT mutants were assessed by western blotting as previously established with minor changes^13^, In detail, 0.5×10^6^ HEK293 cells were seeded onto 6-well plates, transiently transfected the next day with the indicated constructs, and maintained as described above for additional 48h. Cells were then washed with 1 mL of ice-cold PBS and harvested in 150 μL of lysis buffer (150 mM NaCl, 50 mM Tris pH 8.0, 0.5% sodium deoxycholate, 0.1% sodium dodecyl sulfate (SDS), 10 mM Na_4_P_2_O_7_, 1% IGEPAL and 5 mM EDTA) supplemented with a protease inhibitor cocktail (catalog #P8340, Sigma-Aldrich). After incubation in lysis buffer for 60 min at 4°C, lysates were centrifuged for 20 min at 17,000 x *g* at 4°C. Protein concentration in the supernatants was assessed using the *DC^TM^ Protein Assay kit* (Bio-Rad Laboratories) and adjusted to be the same in each condition. Samples were incubated in SDS loading buffer for 60 min at 37°C prior to being resolved by SDS-PAGE and transferred to PVDF membranes. Immunoblotting was carried out using the following antibodies: rat monoclonal anti-DAT (catalog #MAB369, Sigma-Aldrich; dilution 1:1000), rabbit polyclonal anti-DAT raised against the second extracellular loop (catalog #AB5802, Sigma-Aldrich; dilution 1:1000) and HRP-conjugated goat anti-rat IgG antibody (catalog #31470, ThermoFisher; dilution 1:10,000). After detection of DAT from lysates, the membranes were stripped using NaOH 0.4M, then blocked again, and finally probed with an HRP-conjugated monoclonal anti-β-actin antibody (catalog #A3854, Sigma-Aldrich; dilution 1:20,000).

### Immunostaining and confocal microscopy imaging

HEK293 cells transfected with either DAT WT or selected DAT coding variants were seeded onto 96-well black plates (catalog#6005182, PerkinElmer) and maintained as described above. After 48h, cells were fixed for 20 min with a solution containing 3% glyoxal (Sigma-Aldrich), 0.8% acetic acid, adjusted to pH 4.0, then incubated with a blocking buffer containing 5% (v/v) goat serum, 1% (w/v) BSA, for 20 min at room temperature. To detect intracellular and plasma membrane DAT content (total DAT expression), cells were permeabilized with 0.5 % Triton X-100 in PBS for 20 min, and stained applying the monoclonal primary antibody mAB369 (catalog #MAB369, rat, anti-DAT, Sigma-Aldrich; 1:1000). SC-32259 (catalog #sc-32259; rat, anti-DAT; Santa Cruz Biotechnology; 1:200) was applied to non-permeabilized cells, to visualize plasma membrane-expressed DAT. Following two washes in blocking buffer, Alexa488-conjugated secondary antibody (goat anti-rat; Invitrogen; 1:750) was applied both to permeabilized and non-permeabilized cells (60 min at room temperature). After washing and equilibrating with PBS, nuclei were stained with Hoechst33342 (Thermo Fisher Scientific; 1:2000) for 10 min, then maintained in PBS. Confocal images acquisition was carried out on the ImageXpress Confocal HT.ai, applying 20x magnification on 10 randomly selected spots per sample. Images were processed using Fiji-ImageJ 2.14.

### GRAB_DA2M_-sniffer cells assay

GRAB_DA2M_ sniffer cells assay was performed as previously described ^48^. Briefly, sniffer cells were transfected with DAT WT or single DAT mutants and seeded on 96-well plates (catalog#6005689, PerkinElmer Life Science) at a density of ∼3×10^4^ cells/well. Expression of GRAB_DA2M_ sensor was induced by adding 1 µg/mL tetracycline to cell medium. Two days later, cells were washed and equilibrated with aCSF for 5 min, then loaded with 1 µM DA for 15 min. To remove residual extracellular DA, cells were washed twice with aCSF for three minutes, and subsequently buffer was added to let the cells equilibrate for 20 more minutes. To assess AMPH-mediated DA efflux, 30 µM AMPH was administered to each well and fluorescence was measured for 15 minutes. To visualize constitutive DA efflux, cells were washed and equilibrated with aCSF for 5 min, then loaded with 10 µM DA and left for 15 min. After three subsequent washing steps, cells were treated with 10 µM nomifensine and fluorescence was measured for 15 min. Fluorescence intensity (485/520) was recorded every minute throughout the assay, on a POLARstar OMEGA plate reader, at 37 °C.

### Superfusion release assay

Release assays were performed using a superfusion assay as described in detail previously^67^. One day prior to experiments, 100,000 cells expressing the WT or a mutant DAT were seeded in poly-D-lysine (PDL)-coated ibidi µ-slides. On the day of the experiment, cells were preloaded with 0.05 µM [^3^H]MPP^+^ in Krebs-HEPES-buffer (KHB; 120 mM NaCl, 3 mM KCl, 2 mM CaCl_2_·2H_2_O, 2 mM MgCl_2_·6H_2_O, 20 mM D-glucose, pH adjusted to 7.3 with NaOH) for 60 min at 37°C. Cells were then washed once manually with KHB and subsequently superfused with KHB at a rate of 0.5 mL/min for 10 minutes. Basal release was collected in three two-minute fractions in counting vials containing 2 mL scintillation cocktail (Rotiszint Routine, Roth). Subsequently, cells were superfused with 10 µM S-(+)-amphetamine or 10 µM nomifensine in KHB for five fractions. The remaining radioactivity in cells was determined by superfusing cells with 1% SDS for three additional fractions. Radioactivity was measured in a beta-counter (PerkinElmer, Tri-Carb 2810 TR). Release was expressed as percentage of released radioactivity relative to the total radioactivity present at the start of the respective fraction. Experiments were performed in triplicate in five to seven independent experiments.

### Coarse-grained simulations

In absence of the crystal or cryo-EM structure of hDAT in the outward-open conformation at the time of present study, we used Modeller to generate homology models of hDAT based on hSERT structure (PDB ID: 5I71)^68^. To obtain an equilibrated protein-lipid system, the hDAT homology model was simulated in a symmetric membrane composed of 70% 1-palmitoyl-2-oleoyl-sn-glycero-3-phosphocholine (POPC) and 30% cholesterol (CHOL) in a coarse-grained representation for 1 μs. First, the protein was converted to a coarse-grained representation applying Martini forcefield using the martinize.py script^69–71^. After, the lipids were randomly placed around the protein using the insane.py script. The box size used was 10.7 x 10.7 x 10.4 nm. The protein-lipid system was then solvated, neutralized, and NaCl was added to a final concentration of 150 mM. The simulations were run for a total of 1 μs with a time step of 20 fs. Neighbor searching was performed every 20 steps using the Verlet cutoff scheme. The Reaction-Field algorithm was used for electrostatic interactions with a cutoff of 1.1 nm. A single cut-off of 1.1 nm was used for van der Waals interactions. Temperature coupling was done with the V-rescale algorithm and pressure coupling was done with the Parrinello-Rahman algorithm. During the production run, position restraints were applied to the backbone and sidechain atoms with a force constant of 10. Reference coordinates for position restraints were scaled with the pressure coupling. Five independent systems were run.

### All-atoms simulations

The final frame containing protein, membrane, water, and ions of the 1 μs coarse-grained simulation was backmapped to an all-atom representation using the backward.py script^69^. The backmapped protein was then replaced by the original homology model and inserted into the membrane using membed procedure which avoided reducing clashes between the protein and the membrane. As a result, the starting structures of the all-atom simulations consist of the original homology model embedded in an equilibrated lipid environment. Five independent all atom simulations were carried out with GROMACS version 2019.23 applying amber99sb-ildn force field^70,71^. After initial energy minimization, a four-step equilibration protocol of 2.5 ns each was applied to slowly release the position restraints (1000, 100, 10, 1 kJ/mol/nm) applied to the Cα atoms bound ions and dopamine (if present)^72^. The production runs were carried out in absence of any position restraints using a NPT ensemble at 310 K and 1 bar (under semi-isotropic condition). Temperature coupling was done with the V-rescale algorithm and pressure coupling was done with the Parrinello-Rahman algorithm with time constant (tau) set to 0.5 and 20.1 ps, respectively. The van der Waals interactions were described using the Lennard Jones potentials with a cut-off of 0.9 nm. In addition, long-range correction for energy and pressure were applied. The coordinates were saved every 25 ps.

The same starting systems were used to obtain the variants Q317H, S376I, V363I, and S460F and the same simulation parameters were applied. Mutants were created using the PyMOL Mutagenesis Wizard tool^73^.

### Statistical analysis

Data analysis, statistical tests and data plotting were run using GraphPad Prism 10.1.1 (GraphPad Software, San Diego, CA, USA). log*K*_m_ and *V*_max_ values for tritiated DA uptake were calculated using a nonlinear regression analysis with Michaelis-Menten enzyme-kinetics equation Y=*V*_max_*X/(*K*_m_+X). log*K*_d_ and *B*_max_ from DG3-80 saturation binding were obtained using a one-site, specific binding nonlinear regression analysis of baseline-subtracted data, adopting the equation Y=*B*_max_*X/(*K_d_*+X). log*K*_m_DA, log*K*_d_DG3-80, cell-surface DAT expression, total DAT expression, surface/total DAT, *K*_cat_DA, AMPH-induced DA efflux and AMPH sensitivity, constitutive DA efflux, extracellular (upon-nomifensine and baseline) tritiated MPP^+^, and AMPH-induced tritiated MPP^+^ efflux, were statistically examined by applying one-way ANOVA with Holm-Šídák’ multiple comparisons test, comparing each variant to the WT.

## Data availability

The data presented in this study are available from the corresponding author upon request.

## Acknowledgements

This work was supported by the European Union’s Horizon 2020 research and innovation program under the Marie Skłodowska-Curie grant agreement No: 860954 (A.C., U.G.), by the Lundbeck Foundation grants R276-2018-792 (U.G.), R380-2021-1360 (J.D.) and R359-2020-2301 (U.G.) and by the National Institute on Drug Abuse – Intramural Research Program, NIH, Z1A DA000389 (A.H.N.). High content imaging and analysis was performed using the ImageXpress Confocal HT.ai and MetaXpress software (Molecular Devices, Sunnyvale, USA) in the High Content CRISPR Screens (funded by the NNF-0061734) at Biotech Research and Innovation Centre at University of Copenhagen.

## Author contributions

A.C., C.K.H., F.H. and U.G. conceived and administered the project; C.K.H. and U.G. supervised; A.C., F.H. and T.W. carried out genetic data acquisition, management and validation; J.F.S. coordinated ordering of the original DAT mutant construct library and performed initial genotype–phenotype analysis of the variants; A.C. coordinated ordering and validation of a subset of DAT mutant plasmids, performed uptake assays, binding assays, confocal imaging, determination of AMPH response and measurement of constitutive DA efflux; A.H.N. provided the fluorescently-tagged cocaine analogue, DG3-80, synthesized by Dr. Daryl Guthrie in the Medicinal Chemistry Section, NIDA-IRP; J.D. and L.P. performed ELISA and western-blotting; L.A.D.S: and T.S. carried out MD simulations; N.K and H.H.S. provided superfusion release assay data; A.C. and U.G. curated formal data analysis; A.C. and L.A.D.S. designed and prepared the figures; A.C. wrote the first draft, and U.G. wrote-revised and edited the manuscript; A.C., J.D., A.H.N. and U.G. acquired funding; U.G. provided resources. All authors have reviewed the manuscript and agreed to its final version.

## Competing Interests

The authors declare no competing financial interests. The contributions of the NIH author are considered Works of the United States Government. The findings and conclusions presented in this paper are those of the authors and do not necessarily reflect the views of the NIH or the U.S. Department of Health and Human Services.

## Supplementary information

**Supplementary Fig. 1.**
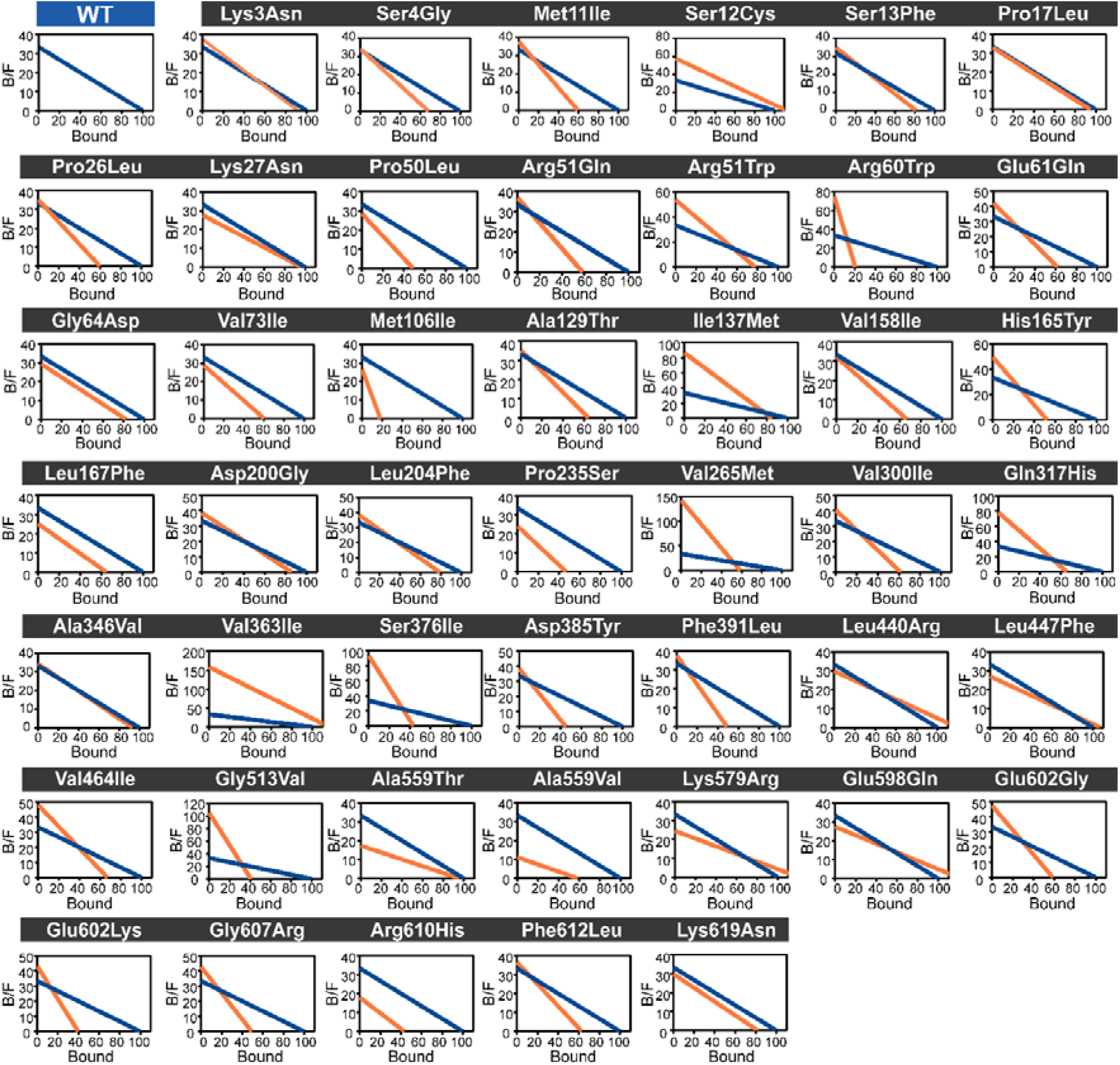
Scatchard plots of cocaine analogue DG3-80 binding to DAT variants. Scatchard plots of fluorescently conjugated cocaine analogue DG3-80 obtained from transiently transfected HEK293 cells expressing either WT DAT (blue) or individual DAT coding variants (orange). Scatchard plots are not shown for variants for which a DG3-80 binding curve could not be fitted. The Scatchard plots are based on values obtained from binding curves fitted using a one-site specific binding nonlinear regression analysis on baseline-subtracted data. B = bound; F = free.

**Supplementary Fig. 2.**
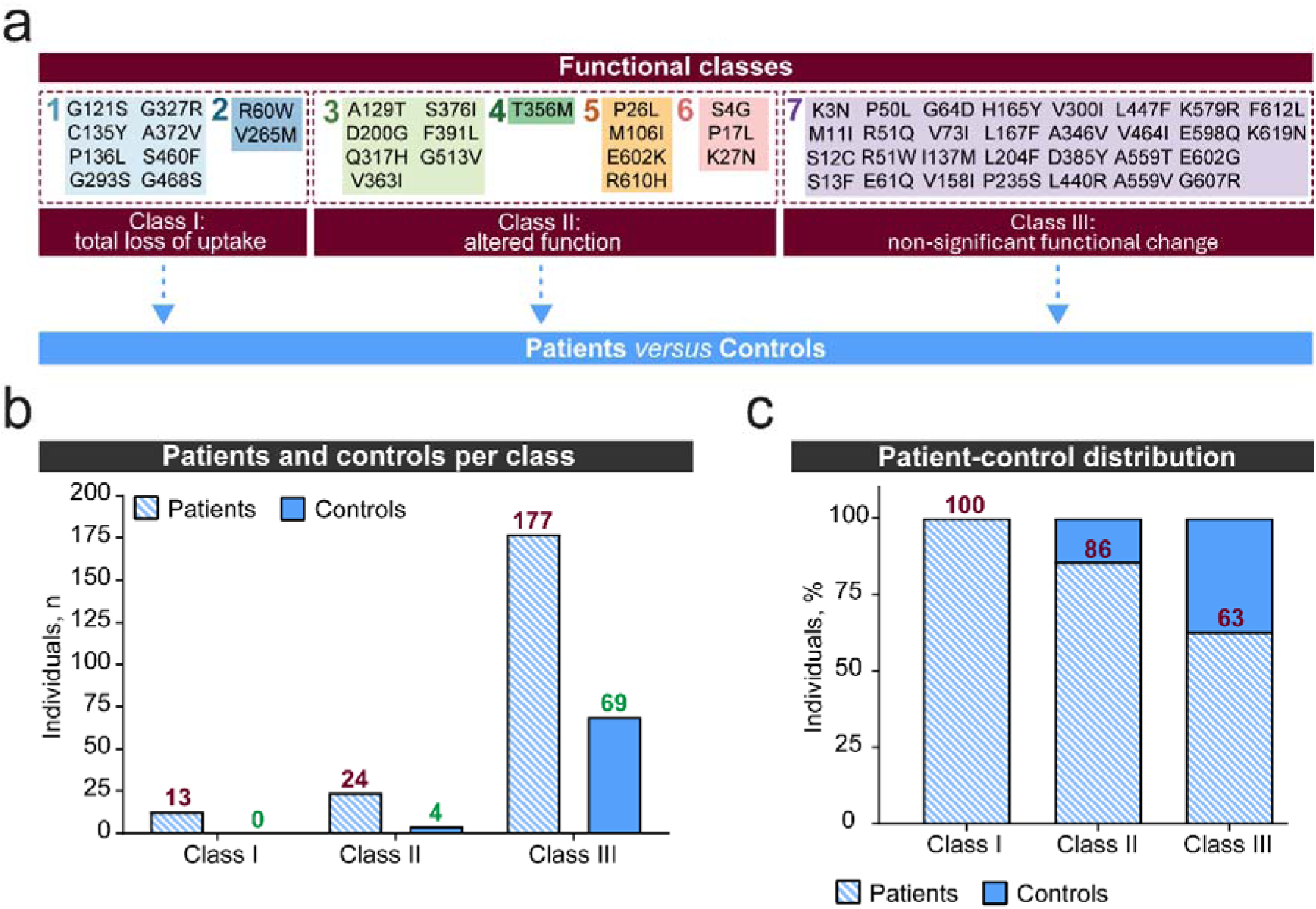
Distribution of patients and controls across functional classes of DAT variants. **a** Graphical overview of DAT variant functional groups categorized into functional classes, and subsequent assessment of patients vs. controls within each class. **b** Bar graph displaying the number of individuals (patients and controls) present in each functional class. **c** Relative distribution (%) of patients and controls across functional classes of DAT variants.

**Supplementary Fig. 3.**
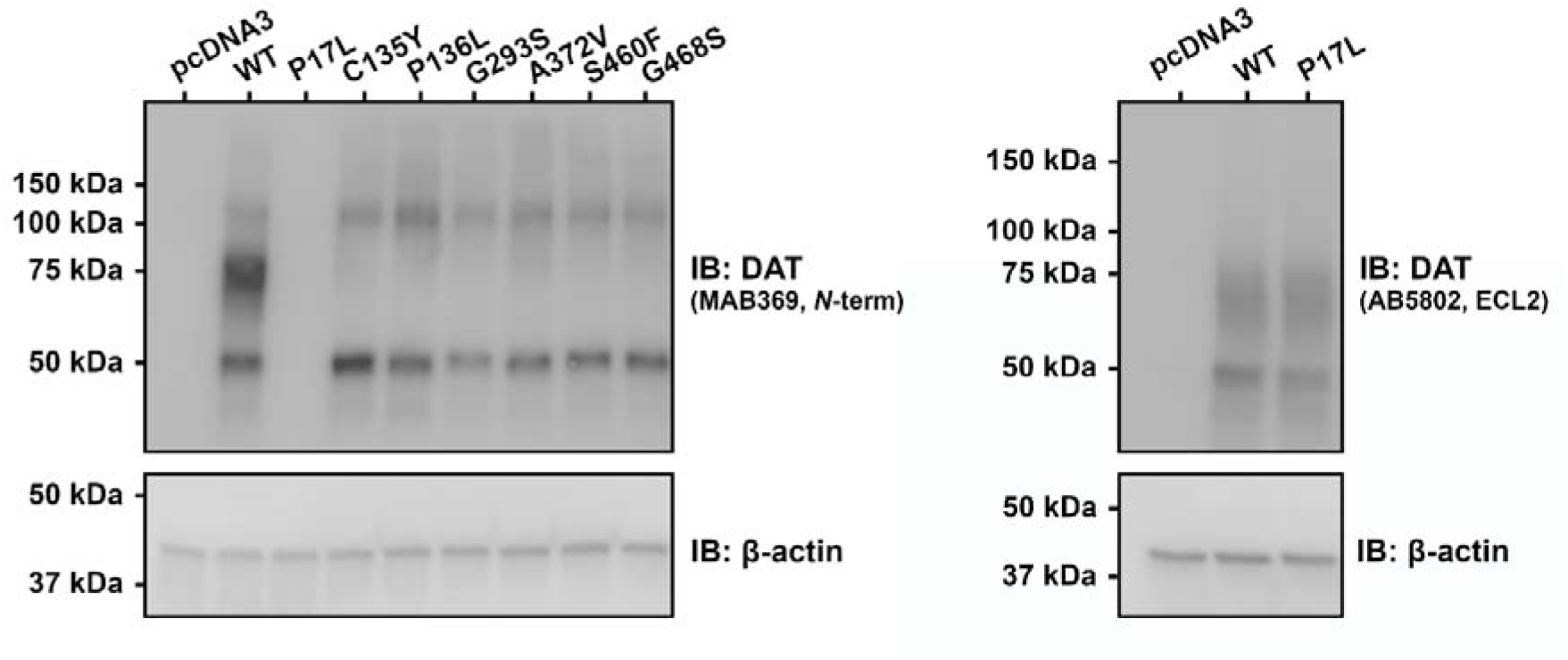
Assessment of expression levels of mutants P17L, C135Y, P136L, G293S, A372V, S460F and G468S through western blotting. **a.** DAT mutants displaying both absent kinetic activity and impaired DG3-80 binding, alongside highly reduced to non-detectable surface and total expression in ELISA (C135Y, P136L, G293S, A372V, S460F and G468S) were blotted using MAB369, which targets DAT N-terminal. P17L was also sampled with western blotting, due to its barely detectable total expression assessed by ELISA. As P17L sits within the MAB369 epitope, no expression is detected. HEK293 cells transfected with either empty pcDNA3 or WT were run as control and reference, respectively, while β-actin was used as loading control. **b.** Expression pattern of P17L tested using AB5802, which targets ECL2, compared to pcDNA3 and WT. β-actin was used as loading control. IB: immunoblotting

**Supplementary Fig. 4.**
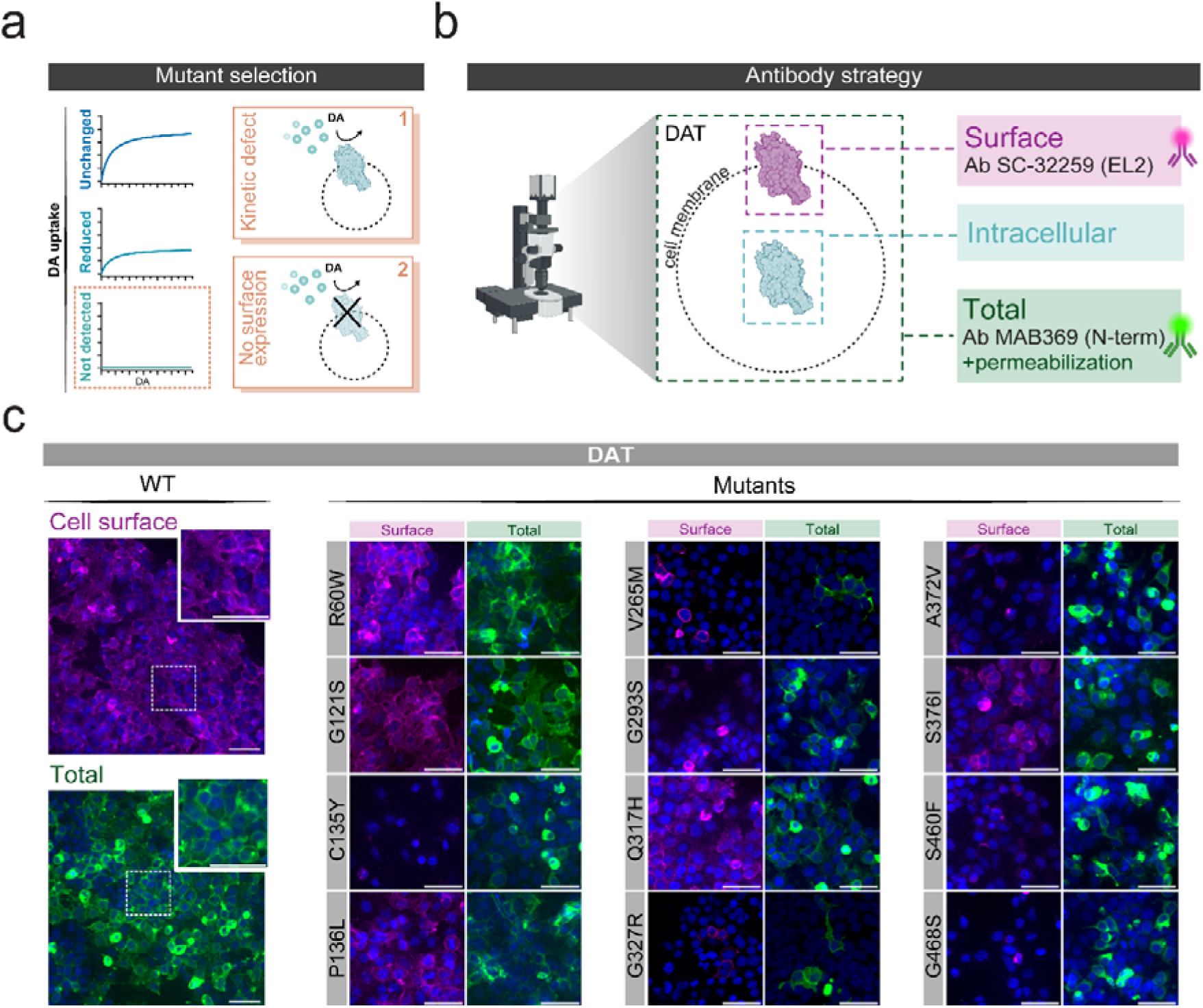
Cell surface expression and total expression levels of highly impaired DAT mutants. **a** Representation of the selection criteria adopted to choose DAT mutants for confocal imaging. Complete ablation of DAT-mediated DA uptake can result from either kinetic defects (box 1) or from complete loss of DAT expression at the cell membrane (box 2). **b** Schematic overview of immunostaining used for confocal microscopy. For detection of plasma membrane DAT (magenta), cells were stained with a primary antibody targeting a portion of DAT ECL2 (Ab: sc-32259). To visualize both surface and cytosolic DAT (green), cells were incubated with a primary antibody recognizing a portion of DAT N-terminal (Ab: MAB369); in this case, cells were permeabilized prior to the staining. **c** Confocal images of HEK293 cells expressing either DAT WT (left), or DAT variants with highly impaired DA uptake (right). For each variant, surface and total DAT expression images are shown side by side. Nuclei were stained using Hoechst 33342. Scale bar = 50 µm.

**Supplementary Fig. 5.**
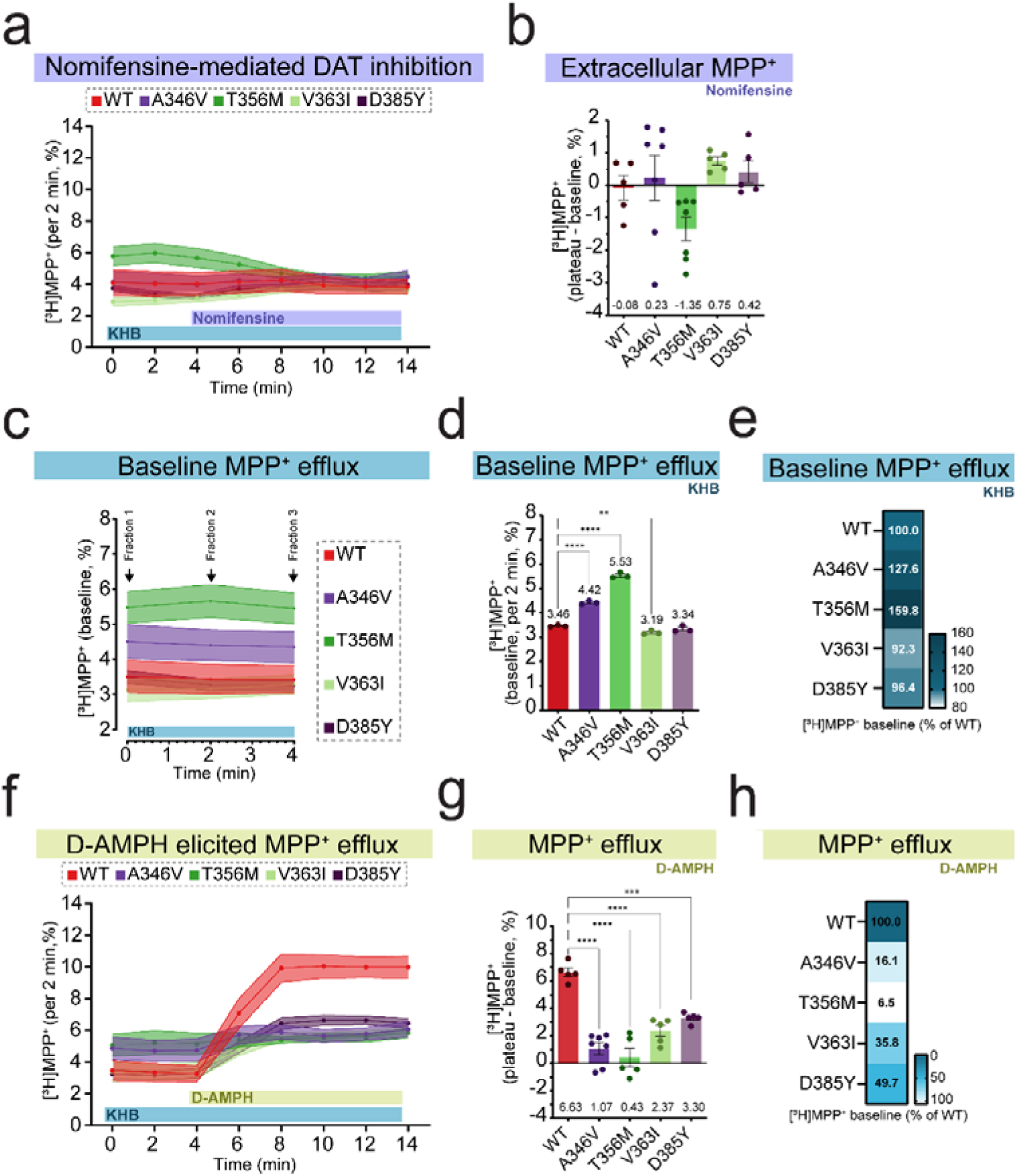
Assessment of constitutive MPP^+^ efflux and AMPH-induced MPP^+^ efflux via DAT on WT, A346V, T356M, V363I and D385Y. **a** Nomifensine inhibition of DAT-mediated MPP^+^ uptake measured on HEK293 cells stably expressing WT or selected DAT mutants, pre-loaded with 0.05 µM [^3^H]MPP^+^ for 60 minutes, to evaluate DAT-mediated constitutive DA efflux. The graph shows curves starting 4 minutes prior to addition of 10 µM nomifensine, and refers to three to seven independent experiments, each performed in duplicate, and normalized to total initial [^3^H]MPP^+^. **b** Extracellular [^3^H]MPP^+^ calculated by baseline (minutes 0-4) subtraction of plateau (minutes 8-14) values reached upon 10 µM nomifensine addition. Data are expressed as a percentage of the total initial [^3^H]MPP^+^. One-way ANOVA with Holm-Šídák’ multiple comparisons test (ns, p > 0.05). **c** Basal [^3^H]MPP^+^ levels measured by superfusing DAT (WT or mutant)-expressing cells with KHB and collecting three fractions (minutes 0, 2 and 4). Data are normalized to the total initial [^3^H]MPP^+^. **d** Quantification of basal [^3^H]MPP^+^ levels of WT and DAT mutants. One-way ANOVA with Holm-Šídák’ multiple comparisons test (**, p < 0.01; ****, p < 0.0001). **e** Heatmap showing [^3^H]MPP^+^ efflux expressed as a percentage of WT basal levels, calculated from data in panel **d**. **f** D-AMPH-elicited [^3^H]MPP^+^ efflux measured on HEK293 cells stably expressing WT or selected DAT mutants, pre-loaded with 0.05 µM [^3^H]MPP^+^ for 60 minutes, Traces start 4 minutes before D-AMPH (10 µM) addition. Values are normalized to total initial [^3^H]MPP^+^ and result from three to seven independent experiments, each performed in duplicate. **g** Net [^3^H]MPP^+^ efflux in response to D-AMPH, obtained by subtracting baseline values (minutes 0-4) from plateau values (minutes 8-14) and expressed as a percentage of the total initial [^3^H]MPP^+^. One-way ANOVA with Holm-Šídák’ multiple comparisons test (***, p < 0.001; ****, p < 0.0001). **h** Heatmap displaying [^3^H]MPP^+^ efflux expressed as a percentage of WT efflux, calculated from values in panel **g**. KHB = Krebs-HEPES-buffer.

**Supplementary Table 1.** Cell surface expression, total expression and catalytic activity of DAT variants.

| DAT variant | DAT expression (E) |  |  | DAT transport rate |
|---|---|---|---|---|
|  | Cell surface (E<sub>CS</sub>, ± S.E.M., % of WT) | Total (E<sub>T</sub>, ± S.E.M., % of WT) | E<sub>CS</sub>,/E<sub>T</sub> (± S.E.M., % of WT) | *K*<sub>cat</sub>DA (± S.E.M., % of WT) |
| WT | 100.0 | 100.0 | 100.0 | 100 |
| Lys3Asn | 92.5 ± 6.8 | 146.8 ± 10.0 | 63.0 ± 6.3 | 155 ± 34 |
| Ser4Gly | 82.0 ± 8.23 | 142.6 ± 13.4 | 57.5 ± 7.9 | 212 ± 58 |
| Met11Ile | 58.5 ± 6.7<sup>**</sup> | 38.9 ± 6.9<sup>**</sup> | 150.3 ± 31.6 | 121 ± 21 |
| Ser12Cys | 96.9 ± 1.1 | 59.8 ± 4.5 | 162.0 ± 12.4 | 81 ± 7 |
| Ser13Phe | 71.2 ± 9.4 | 70.4 ± 11.9 | 101.2 ± 21.7 | 218 ± 35 |
| Pro17Leu | 74.8 ± 8.1 | Not detected | - | 293 ± 104<sup>***</sup> |
| Pro26Leu | 80.0 ± 2.2 | 90.4 ± 2.8 | 88.5 ± 3.7 | 65 ± 9 |
| Lys27Asn | 81.0 ± 8.0 | 142.3 ± 16.6 | 56.9 ± 8.7 | 324 ± 118<sup>***</sup> |
| Pro50Leu | 79.8 ± 6.8 | 112.3 ± 13.2 | 71.1 ± 10.3 | 195 ± 38 |
| Arg51Gln | 90.7 ± 1.7 | 77.08 ± 12.7 | 113.2 ± 9.3 | 148 ± 30 |
| Arg51Trp | 91.8 ± 3.9 | 104.0 ± 7.3 | 108.6 ± 5.8 | 116 ± 21 |
| Arg60Trp | 40.1 ± 6.4<sup>****</sup> | 73.5 ± 11.3 | 54.6 ± 12.1 | - |
| Glu61Gln | 81.7 ± 7.0 | 117.6 ± 14.0 | 69.5 ± 10.2 | 113 ± 23 |
| Gly64Asp | 76.8 ± 8.4 | 112.2 ± 11.1 | 68.4 ± 10.1 | 164 ± 26 |
| Val73Ile | 70.6 ± 11.6 | 115.7 ± 17.5 | 61.0 ± 13.6 | 185 ± 25 |
| Met106Ile | 48.7 ± 7.4<sup>****</sup> | 89.8 ± 14.7 | 54.3 ± 12.1 | 134 ± 14 |
| Gly121Ser | 54.8 ± 10.2<sup>***</sup> | 81.3 ± 11.3 | 67.5 ± 15.6 | - |
| Ala129Thr | 82.3 ± 2.1 | 113.2 ± 1.5 | 72.7 ± 2.1 | 55 ± 7 |
| Cys135Tyr | Not detected | Not detected | - | - |
| Pro136Leu | Not detected | Not detected | - | - |
| Ile137Met | 74.7 ± 9.8 | 98.4 ± 12.6 | 75.9 ± 13.9 | 199 ± 57 |
| Val158Ile | 63.2 ± 11.1<sup>*</sup> | 92.2 ± 14.4 | 68.6 ± 16.1 | 162 ± 16 |
| His165Tyr | 52.2 ± 8.5<sup>***</sup> | 84.0 ± 13.8 | 62.1 ± 14.3 | 124 ± 28 |
| Leu167Phe | 62.6 ± 9.4<sup>**</sup> | 97.3 ± 18.2 | 64.4 ± 15.4 | 128 ± 30 |
| Asp200Gly | 58.2 ± 9.4<sup>**</sup> | 108.6 ± 14.5 | 53.6 ± 11.2 | 63 ± 10 |
| Leu204Phe | 65.2 ± 9.8<sup>*</sup> | 119.6 ± 17.3 | 54.6 ± 11.4 | 145 ± 17 |
| Pro235Ser | 70.7 ± 9.9 | 119.2 ± 15.6 | 59.3 ± 11.4 | 121 ± 15 |
| Val265Met | 70.4 ± 2.3 | 97.4 ± 2.1 | 72.3 ± 2.8 | - |
| Gly293Ser | Not detected | Not detected | - | - |
| Val300Ile | 82.5 ± 14.9 | 98.3 ± 12.1 | 84.0 ± 18.4 | 114 ± 37 |
| Gln317His | 41.6 ± 8.3<sup>****</sup> | 68.6 ± 11.4 | 60.6 ± 15.8 | - |
| Gly327Arg | 27.1 ± 5.2<sup>****</sup> | 44.7 ± 6.5<sup>*</sup> | 60.7 ± 14.6 | - |
| Ala346Val | 73.2 ± 12.6 | 81.4 ± 9.2 | 89.9 ± 18.5 | 95 ± 19 |
| Thr356Met | 60.4 ± 8.5<sup>**</sup> | 91.5 ± 15.8 | 66.0 ± 14.7 | 70 ± 12 |
| Val363Ile | 68.8 ± 12.1 | 111.4 ± 20.2 | 61.8 ± 15.6 | 30 ± 5 |
| Ala372Val | Not detected | 15.2 ± 4.3<sup>****</sup> | - | - |
| Ser376Ile | 10.7 ± 2.4<sup>****</sup> | 30.7 ± 8.2<sup>**</sup> | 34.7 ± 12.1<sup>*</sup> | 151 ± 57 |
| Asp385Tyr | 67.5 ± 2.1<sup>*</sup> | 81.4 ± 1.7 | 82.8 ± 3.1 | 123 ± 31 |
| Phe391Leu | 26.6 ± 6.2<sup>****</sup> | 50.5 ± 11.4 | 52.6 ± 17.0 | 112 ± 20 |
| Leu440Arg | 68.2 ± 15.4<sup>*</sup> | 100.2 ± 19.3 | 68.0 ± 20.2 | 143 ± 18 |
| Leu447Phe | 72.6 ± 13.8 | 100.0 ± 19.5 | 72.6 ± 19.8 | 76 ± 12 |
| Ser460Phe | Not detected | Not detected | - | - |
| Val464Ile | 77.2 ± 2.5 | 106.0 ± 5.0 | 72.9 ± 4.1 | 87 ± 9 |
| Gly468Ser | Not detected | Not detected | - | - |
| Gly513Val | 42.1 ± 8.8<sup>****</sup> | 79.8 ± 16.9 | 52.8 ± 15.7 | 40 ± 11 |
| Ala559Thr | 68.1 ± 13.5<sup>*</sup> | 109.6 ± 19.2 | 62.1 ± 16.4 | 114 ± 16 |
| Ala559Val | 64.9 ± 14.0<sup>*</sup> | 108.1 ± 16.6 | 60.0 ± 15.9 | 131 ± 14 |
| Lys579Arg | 63.1 ± 11.6<sup>*</sup> | 110.8 ± 18.9 | 56.9 ± 14.3 | 129 ± 20 |
| Glu598Gln | 70.2 ± 13.3 | 110.0 ± 23.5 | 63.8 ± 18.2 | 138 ± 22 |
| Glu602Gly | 51.2 ± 6.7<sup>**</sup> | 110.2 ± 17.4 | 46.5 ± 9.5 | 282 ± 41<sup>**</sup> |
| Glu602Lys | 56.9 ± 6.4<sup>**</sup> | 103.3 ± 14.9 | 55.1 ± 10.1 | 142 ± 22 |
| Gly607Arg | 46.5 ± 5.6<sup>****</sup> | 81.4 ± 12.8 | 57.2 ± 11.3 | 122 ± 18 |
| Arg610His | 54.6 ± 8.8<sup>***</sup> | 106.8 ± 17.9 | 51.1 ± 11.9 | 149 ± 9 |
| Phe612Leu | 56.0 ± 9.3<sup>***</sup> | 103.8 ± 22.1 | 54.0 ± 14.6 | 239 ± 88 |
| Lys619Asn | 54.4 ± 9.8<sup>***</sup> | 113.6 ± 21.1 | 47.9 ± 12.4 | 138 ± 20 |

Values indicating cell surface (E<sub>CS</sub>) and total expression (E<sub>T</sub>) of each DAT variant, as percentage of WT, assessed by ELISA upon transient transfection of HEK293 cells with either WT or DAT mutants. Transport rate of each DAT mutant calculated as *K*<sub>cat</sub>DA = *V*<sub>max</sub>DA/ E<sub>CS</sub>, expressed as percentage of WT. Data are presented as mean ± S.E.M. of five independent experiments, each performed in triplicates (cell surface expression) or duplicate (total expression). One-way ANOVA with Holm-Šídák' multiple comparisons test (\*, p < 0.05; \*\*, p < 0.01; \*\*\*, p < 0.001; \*\*\*\*, p < 0.0001).

**Supplementary Table 2.**
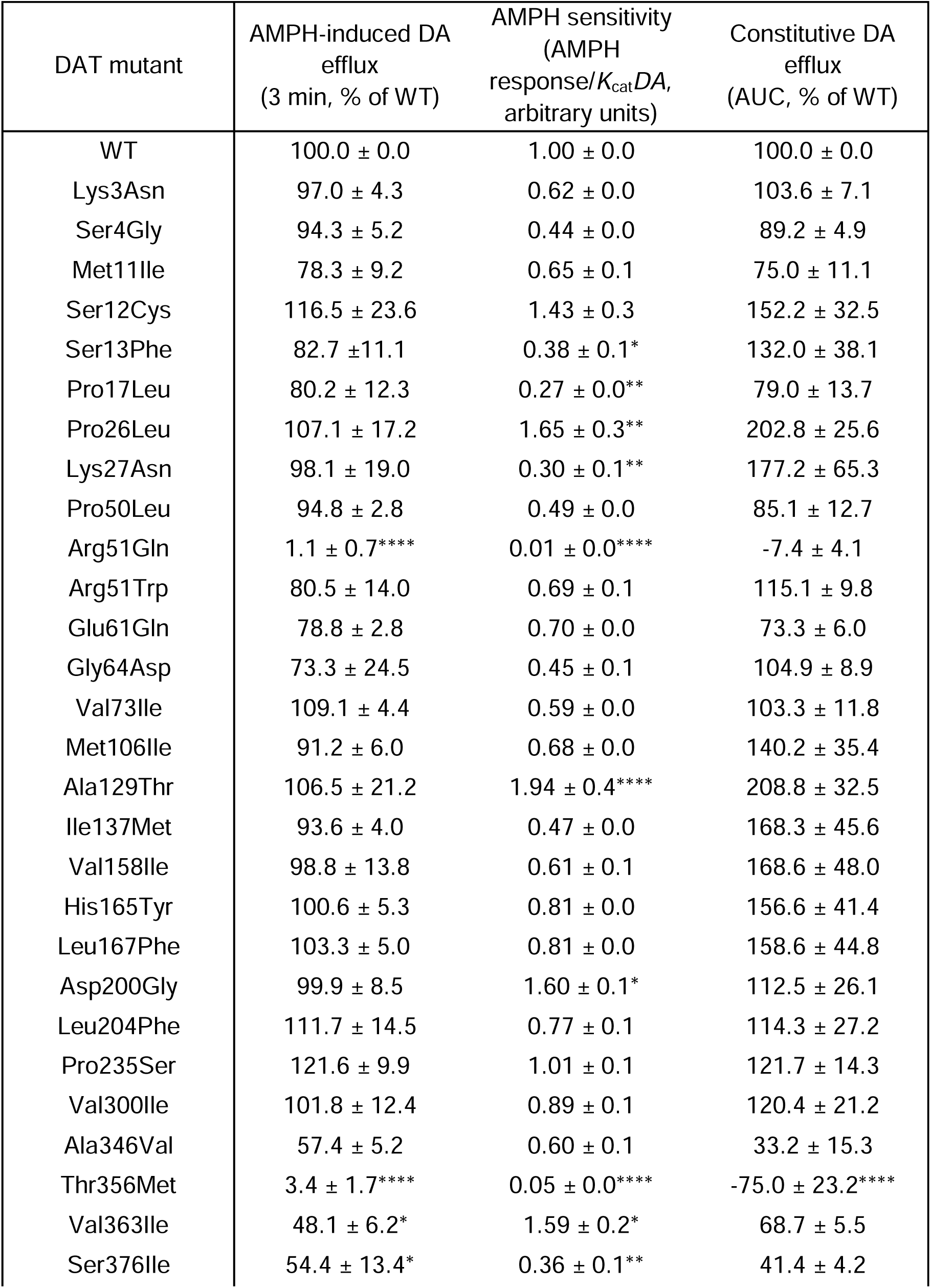
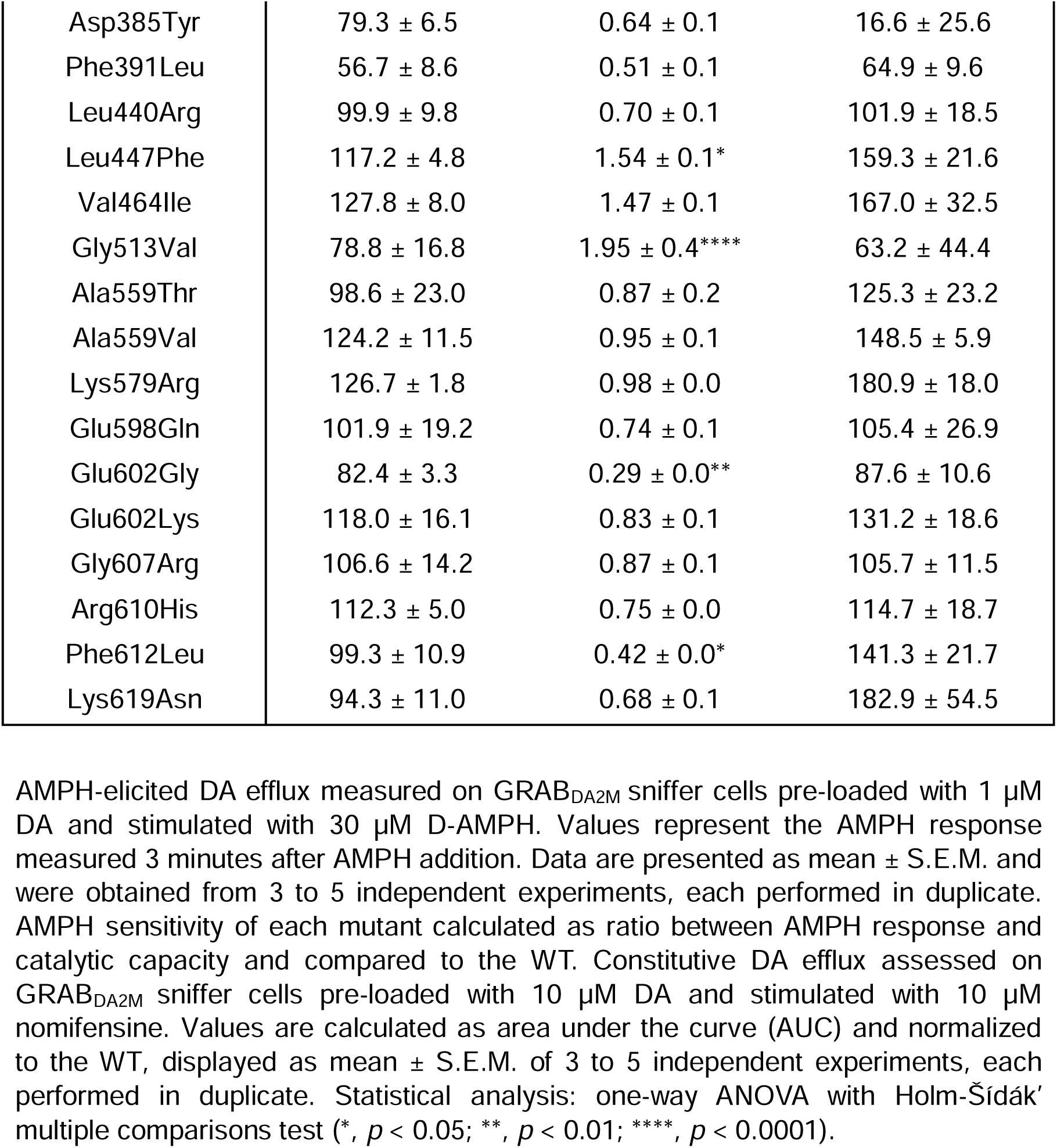
AMPH-induced DA efflux and constitutive DA efflux assessed on GRAB_DA2M_ sniffer cells.

| DAT mutant | AMPH-induced DA efflux<br>(3 min, % of WT) | AMPH sensitivity<br>(AMPH response/*K*<sub>cat</sub>*DA*, arbitrary units) | Constitutive DA efflux<br>(AUC, % of WT) |
|---|---|---|---|
| WT | 100.0 ± 0.0 | 1.00 ± 0.0 | 100.0 ± 0.0 |
| Lys3Asn | 97.0 ± 4.3 | 0.62 ± 0.0 | 103.6 ± 7.1 |
| Ser4Gly | 94.3 ± 5.2 | 0.44 ± 0.0 | 89.2 ± 4.9 |
| Met11Ile | 78.3 ± 9.2 | 0.65 ± 0.1 | 75.0 ± 11.1 |
| Ser12Cys | 116.5 ± 23.6 | 1.43 ± 0.3 | 152.2 ± 32.5 |
| Ser13Phe | 82.7 ±11.1 | 0.38 ± 0.1* | 132.0 ± 38.1 |
| Pro17Leu | 80.2 ± 12.3 | 0.27 ± 0.0** | 79.0 ± 13.7 |
| Pro26Leu | 107.1 ± 17.2 | 1.65 ± 0.3** | 202.8 ± 25.6 |
| Lys27Asn | 98.1 ± 19.0 | 0.30 ± 0.1** | 177.2 ± 65.3 |
| Pro50Leu | 94.8 ± 2.8 | 0.49 ± 0.0 | 85.1 ± 12.7 |
| Arg51Gln | 1.1 ± 0.7**** | 0.01 ± 0.0**** | -7.4 ± 4.1 |
| Arg51Trp | 80.5 ± 14.0 | 0.69 ± 0.1 | 115.1 ± 9.8 |
| Glu61Gln | 78.8 ± 2.8 | 0.70 ± 0.0 | 73.3 ± 6.0 |
| Gly64Asp | 73.3 ± 24.5 | 0.45 ± 0.1 | 104.9 ± 8.9 |
| Val73Ile | 109.1 ± 4.4 | 0.59 ± 0.0 | 103.3 ± 11.8 |
| Met106Ile | 91.2 ± 6.0 | 0.68 ± 0.0 | 140.2 ± 35.4 |
| Ala129Thr | 106.5 ± 21.2 | 1.94 ± 0.4**** | 208.8 ± 32.5 |
| Ile137Met | 93.6 ± 4.0 | 0.47 ± 0.0 | 168.3 ± 45.6 |
| Val158Ile | 98.8 ± 13.8 | 0.61 ± 0.1 | 168.6 ± 48.0 |
| His165Tyr | 100.6 ± 5.3 | 0.81 ± 0.0 | 156.6 ± 41.4 |
| Leu167Phe | 103.3 ± 5.0 | 0.81 ± 0.0 | 158.6 ± 44.8 |
| Asp200Gly | 99.9 ± 8.5 | 1.60 ± 0.1* | 112.5 ± 26.1 |
| Leu204Phe | 111.7 ± 14.5 | 0.77 ± 0.1 | 114.3 ± 27.2 |
| Pro235Ser | 121.6 ± 9.9 | 1.01 ± 0.1 | 121.7 ± 14.3 |
| Val300Ile | 101.8 ± 12.4 | 0.89 ± 0.1 | 120.4 ± 21.2 |
| Ala346Val | 57.4 ± 5.2 | 0.60 ± 0.1 | 33.2 ± 15.3 |
| Thr356Met | 3.4 ± 1.7**** | 0.05 ± 0.0**** | -75.0 ± 23.2**** |
| Val363Ile | 48.1 ± 6.2* | 1.59 ± 0.2* | 68.7 ± 5.5 |
| Ser376Ile | 54.4 ± 13.4* | 0.36 ± 0.1** | 41.4 ± 4.2 |
| Asp385Tyr | 79.3 ± 6.5 | 0.64 ± 0.1 | 16.6 ± 25.6 |
| Phe391Leu | 56.7 ± 8.6 | 0.51 ± 0.1 | 64.9 ± 9.6 |
| Leu440Arg | 99.9 ± 9.8 | 0.70 ± 0.1 | 101.9 ± 18.5 |
| Leu447Phe | 117.2 ± 4.8 | 1.54 ± 0.1* | 159.3 ± 21.6 |
| Val464Ile | 127.8 ± 8.0 | 1.47 ± 0.1 | 167.0 ± 32.5 |
| Gly513Val | 78.8 ± 16.8 | 1.95 ± 0.4**** | 63.2 ± 44.4 |
| Ala559Thr | 98.6 ± 23.0 | 0.87 ± 0.2 | 125.3 ± 23.2 |
| Ala559Val | 124.2 ± 11.5 | 0.95 ± 0.1 | 148.5 ± 5.9 |
| Lys579Arg | 126.7 ± 1.8 | 0.98 ± 0.0 | 180.9 ± 18.0 |
| Glu598Gln | 101.9 ± 19.2 | 0.74 ± 0.1 | 105.4 ± 26.9 |
| Glu602Gly | 82.4 ± 3.3 | 0.29 ± 0.0** | 87.6 ± 10.6 |
| Glu602Lys | 118.0 ± 16.1 | 0.83 ± 0.1 | 131.2 ± 18.6 |
| Gly607Arg | 106.6 ± 14.2 | 0.87 ± 0.1 | 105.7 ± 11.5 |
| Arg610His | 112.3 ± 5.0 | 0.75 ± 0.0 | 114.7 ± 18.7 |
| Phe612Leu | 99.3 ± 10.9 | 0.42 ± 0.0* | 141.3 ± 21.7 |
| Lys619Asn | 94.3 ± 11.0 | 0.68 ± 0.1 | 182.9 ± 54.5 |

AMPH-elicited DA efflux measured on GRAB<sub>DA2M</sub> sniffer cells pre-loaded with 1 µM DA and stimulated with 30 µM D-AMPH. Values represent the AMPH response measured 3 minutes after AMPH addition. Data are presented as mean ± S.E.M. and were obtained from 3 to 5 independent experiments, each performed in duplicate. AMPH sensitivity of each mutant calculated as ratio between AMPH response and catalytic capacity and compared to the WT. Constitutive DA efflux assessed on GRAB<sub>DA2M</sub> sniffer cells pre-loaded with 10 µM DA and stimulated with 10 µM nomifensine. Values are calculated as area under the curve (AUC) and normalized to the WT, displayed as mean ± S.E.M. of 3 to 5 independent experiments, each performed in duplicate. Statistical analysis: one-way ANOVA with Holm-Šídák' multiple comparisons test (\*, *p* < 0.05; \*\*, *p* < 0.01; \*\*\*\*, *p* < 0.0001).

